# Altered bacteria community dominance reduces tolerance to resident fungus and seed to seedling growth performance in maize (*Zea mays* L. var. DBK 177)

**DOI:** 10.1101/2020.07.16.206441

**Authors:** Lidiane Figueiredo dos Santos, Julie Fernandes Souta, Letícia Oliveira da Rocha, Cleiton de Paula Soares, Maria Luiza Carvalho Santos, Clicia Grativol Gaspar de Matos, Luiz Fernando Wurdig Roesch, Fabio Lopes Olivares

## Abstract

Seeds are reservoirs of beneficial and harmful microorganism that modulates plant growth and health. Here, we access seed to seedling bacteriome assembly modified by seed-disinfection and the underlined effect over maize germination performance and root-seedlings microbial colonization. Seed-disinfection was performed with sodium hypochlorite (1.25%, 30 min), resulting in a reduction of the cultivable-dependent fraction of seed-borne bacteria population, but not significantly detected by real-time PCR, microscopy, and biochemical analysis of the roots on germinated seeds. 16S rRNA sequencing revealed that the seed and root bacteriome exhibited similar diversity and did not differ in the structure concerning seed-disinfection. On the other hand, the abundance reduction of the genera f_Enterobacteriaceae_922761 (unassigned genus), *Azospirillum,* and *Acinetobacter* in disinfected-seed prior germination seems to display changes in prominence of several new taxa in the roots of germinated seeds. Interestingly, this reduction in the bacteriome negatively affected the germination speed and growth of maize plantlets. Additionally, bacteriome re-shape increased the maize var DKB 177 susceptible to the seed-borne plant pathogen *Penicillium* sp. Such changes in the natural seed-borne composition removed the natural barrier, increasing susceptibility to pathogens, impairing disinfected seeds to germinate, and develop. We conclude that bacteria borne in seeds modulate the relative abundance of taxa in the root, promote germination, seedling growth, and protect the maize against fungal pathogens.

## Introduction

Plants are colonized by diverse microbial assemblages, known as microbiota (set of microorganisms) or microbiome (set of genomes) (Compant et al., 2019). Regarding the bacterial component of the microbiome (bacteriome), the ability to occupy various niches in the plant (surface and interior of the tissues) (Gopal and Gupta, 2018) and perform various beneficial activities, such as promoting growth and biocontrol of phytopathogens, is highlighted. These bacteria generally promote plant growth by facilitating the acquisition of nutrients (nitrogen, phosphorus, and iron) and producing/modulating phytohormones (auxin, gibberellin, and cytokinin) (direct effect); or by reducing damage caused by fungal and bacterial (harmful) pathogens through compounds they produce (siderophores, antibiotics, lytic enzymes, bacteriocins, lipoproteins, and volatile organic compounds) (indirect effect) (Orozco-Mosqueda et al., 2018, Verma et al., 2019b). Direct and indirect interactions of the microbiota with plants are essential to reduce the use of synthetic and pesticide fertilizers and make agriculture more sustainable.

The plant bacteriome has its origin in seeds (endo and epiphytic), considered a natural carrier of microbial inoculants transmitted vertically (Frank et al., 2017). Bacteria have already been isolated from sterilized seeds on the surface of many cultures (Verma et al., 2017, Verma et al., 2018, Verma and White, 2018), which suggests their protection inside the seed or strong adhesion to the surface. During germination, the radicle, already densely colonized by resident seed-bacteria, elongates and emerges from its coating. Then, the primary root grows in contact with the soil, which becomes a new source of bacteria for the plant host via a horizontal transmission (Bakker et al., 2015). Highlighting only the bacterial associations originating from the seed, we found studies that demonstrated the ability of its prokaryotic inhabitants to promote germination and growth of different plant species, which was confirmed by removing them by chemical and thermal disinfection (Holland, 2016, Irizarry and White, 2017, Verma et al., 2017, Verma et al., 2018, Verma and White, 2018, Holland, 2019, Verma et al., 2019b).

In addition to bacteria, the seeds harbor fungi of a phytopathogenic nature that can develop during the germination period, delaying germination, or killing the seed (Xing et al., 2018). Among the main pathogens transmitted by seed are fungi of the genus *Penicillium*, which infect a wide range of economically important plants (Xing et al., 2018). These fungi often reduce grain yield and quality, in addition to producing mycotoxins (Akonda et al., 2016). Fortunately, seed bacteria are competent biocontrol agents and protect plants from their enemies (Verma et al., 2017, Khalaf and Raizada, 2018, Verma et al., 2018, Verma and White, 2018, White et al., 2017).

Most of the analyzes available on the seed microbiota focus on the diversity or functional abilities of isolated bacteria. In the present study, possible roles for the seed-borne bacteria community were accessed by comparing chemical disinfected and non-disinfected maize seeds growing under axenic conditions. Using cultivable-independent approaches, we evaluated bacteria populations of non-germinated seeds and radicle of germinated seeds throughout 16S rRNA sequencing and population size by real-time PCR. Also, the population size of the cultivable bacteria pool and the structural interaction between the seed-resident microbial community and maize seedling were accessed. Finally, seed germination, seedling growth, and stored reserve remobilization were evaluated, and during these assays, it was observed a differential behavior for the seed-borne fungus *Penicillium* sp. on disinfected and non-disinfected seeds.

## Material and Methods

### Surface disinfection, germination, and seedlings growth promotion of maize

Maize seeds (*Zea mays* L.) of the DKB 177 variety (Dekalb®, Brazil) were immersed in sterile distilled water for 5 h, with part of the seeds being non-disinfected (*NDS treatment*). The disinfected seeds (*DS treatment*) were treated in 70% alcohol for 5 min and sodium hypochlorite (NaClO; Butterfly Ecologia, Audax Company) 1.25% for 30 min. Washings in sterile distilled water were performed between the solutions (1x) and after immersion in hypochlorite (5x). Preliminary tests were carried out to reach the 30 min time of immersion in NaClO (1.25%). For this, seeds were immersed in this solution for times ranging from zero to 90 min (0.10, 20, 30, 40, 50, 60, 70, 80 and 90 min) and placed in tubes containing NB liquid medium (Nutrient Broth; 5 mL) for 48 h (180 rpm, up to 30 ° C). The optical density (OD) of each treatment was measured at 595 nm in a spectrophotometer (Chameleon, Hidex, model 425-156) and associated with the growth of seed bacteria.

The effect of disinfecting seeds for 30 min was evaluated using the Live/Dead® kit (Thermo Fisher Scientific), which analyzes the viability of bacterial cells. For that, bacterial suspension (2 mL) grown from the seed (zero- and 30-min immersion in hypochlorite) was centrifuged (10,000x g for 4 min) and resuspended in saline (NaCl; 8.5 g L-1) twice. Then, two fluorescent nucleic acid dyes, SYTO9 and propidium iodide (PI) (0.5 μL each) were mixed with the bacterial suspension (100 μL) and incubated at room temperature for 15 min. The stained bacteria were visualized using an Axioplan Zeiss epifluorescence microscope.

Disinfected and non-disinfected seeds were germinated under axenic conditions, being placed in Petri dishes (8 repetitions; 12 seeds per dish) containing agar-water medium (0.5%) and packed in BOD (30 °C; photoperiod 12/12 h (light/dark)) for 5 days. After this period, germination percentage (%G), germination speed index (GSI), average germination time (AGT), and average germination speed (AGS) (Maguire, 1962) were evaluated using an unpaired t-test (PRISM, version 8). Radicles ≥ 5 mm were considered to be germinated.

### Seed-borne bacteria population count

The bacteria population associated to emerged radicle of germinated seedlings under axenic conditions (as described in topic 2.1) of disinfected and non-disinfected seeds were determined by Most Probable Number (MPN) technique for positive growth in semi-solid media (data expressed as log_10_ n° cells. g^−1^ root) for diazotrophic population estimation and by the colony-forming unit in Nutrient Broth (NB) solid medium plates (data expressed as log_10_ n° cells. g^−1^ root or mL^−1^) for total bacteria or total bacteria after population enrichment in liquid NB medium. For this, roots (1 g) were macerated in saline (NaCl; 99 mL; 8.5 g L^−1^), subjected to serial dilution (10^−3^ to 10^−6^) and inoculated with a pipette (100 μL) into a 16 mL glass flask containing 5 mL of N-free semi-solid JNFb and LGI medium (respectively containing malic acid and sucrose as C-source) or plated in NB medium. The flasks and Petri plates were incubated in BOD (30 °C; 5 to 7 days), and the counts were as described above and following Baldani et al. (2014). Additionally, primary root segments were placed in tubes containing NB liquid medium (5 mL) subjected to “overnight” agitation (180 rpm, at 30 °C). After bacterial growth, serial dilution, plating in solid NB, and colony counting were performed.

### Structural characterization of the maize bacterial microbiota by light microscopy (LM), scanning electron microscopy (SEM) and transmission electron microscopy (TEM)

Maize seeds and seedlings were grown under axenic conditions were collected and processed for light microscopy (LM) and scanning electron microscopy (SEM).

For LM, the primary maize root was visualized after being stained with 2,3,5-triphenyl tetrazolium chloride (0.1% TTC for 2 h) (TTC; Reagen®), followed by reduction of the tissue background after immersing the root in potassium hydroxide solution (2.5% KOH for 40 min). Stained roots were placed on slides with sterile distilled water and bacteria visualized for reducing the TTC from the colorless soluble form to the insoluble pink form, with this precipitation around the colonies being recorded by the bright-field microscope Axioplan Zeiss.

For SEM, segments (1 cm) of the primary maize root were fixed in glutaraldehyde (2.5%) and paraformaldehyde (4%) in sodium phosphate buffer (0.05 mol L^−1^, pH 7.0). Then, the segments were washed with the same buffer (3 times for 20 min for root; 30 min for seed) and dehydrated in an alcoholic series (15, 30, 50, 70, 90 and 2 × 100% at 15 min for each root; 30 min for seed). The samples were dried in a critical point device (Bal-tec CPD 030), mounted on aluminum stubs, and metalized with ionized platinum in a sputtering coat apparatus (Bal-tec SCD 050). Maize seeds and roots were visualized in SEM Zeiss EVO 40 at 15 kV. Magnifying glass (Zeiss Stemi SV 11). Magnifying glass records (Zeiss Stemi SV 11) were used to indicate macroscopic structures of the seed.

Other root segments were incorporated, after the dehydration phase, in crescent LR White resin (medium grade) until complete tissue replacement of ethanol for resin. After that, individual samples were mounted in transparent gelatin capsules filled with pure fresh resin and then polymerized in an oven at 60 °C for 24 h. Semi-thin sections (0.8-1.0 μm) of cured samples were obtained with the aid of a glass knife and ultramicrotome (Reichert-Jung Ultracut II E). The sections were collected on glass slides heated in a metal plate and stained with toluidine blue (1%). After staining, the material was mounted in water with a coverslip, and it was observed under a light microscope. For TEM, samples prepared as described above were sectioned in ultra-thin sections (50-90 nm) with the aid of a diamond knife and ultramicrotome (Reichert-Jung Ultracut II E). The sections collected in copper grids (300 mesh), contrasted with uranyl acetate (5% for 20 min) and lead citrate (5 min) and observed in a transmission electron microscope JEOL 1400 Plus at 80Kv.

### Maize seeds and roots bacteriome

Seeds and roots were sampled from plates of the axenic assay described above and stored at −70 °C until DNA extraction. Frozen samples were macerated in liquid nitrogen to extract genomic DNA (from 0.2 g) using Cetyltrimethylammonium bromide (CTAB) for roots and seeds (Chen and Ronald, 1999, Doyle and Doyle, 1987). The amount of DNA in the samples was determined using NanoDrop 2000® spectrophotometer (Thermo Scientific) and Qubit® fluorometer (Invitrogen), while the quality was confirmed in agarose gel (0.8%) electrophoresis (80 V, for 70 min). The total DNA was sent to the company “WEMSeq Biotechnology” for sequencing of the 16S rRNA gene in Illumina MiSeq, with three replicates per treatment. The samples were amplified with primers 515F and 806R against the V4 region of the 16S rRNA (Caporaso et al., 2012). PCR products were quantified (Qschd dsDNA HS kit, Invitrogen) and sequenced on the Illumina MiSeq platform (300V2 Kit, Illumina) according to the manufacturer’s instructions.

The MiSeq raw sequences were analyzed in QIIME (Caporaso et al., 2010), version 1.9.0, where low-quality readings were filtered, and the rest grouped into Operational Taxonomic Units (OTUs) using a 97% identity threshold. After grouping, sequences were aligned and classified with the SILVA database (Quast et al., 2012). The quality of the sampling was estimated from Good’s coverage (Good, 1953). Subsequent analyzes were performed in the R environment (Team, 2013) using the phyloseq package (McMurdie and Holmes, 2013) to estimate the alpha and beta diversity. The ordering of the sequencing for beta diversity was performed based on the Bray-Curtis dissimilarity matrix and presented in principal coordinate analysis (PCoA) graphs. Permutational multivariate analysis of variance (Permanova) (Anderson, 2014) was used to assess statistical differences between treatments through the vegan package (Oksanen et al., 2013). The alpha diversity of the treatments was estimated from the Shannon index, and the results were contrasted using the Wilcoxon nonparametric statistical method. Venn diagrams were created to illustrate the overlap of OTUs between samples. Differential and relative abundance analyzes were performed at the gender level, the latter being represented in heatmaps.

### Quantitative PCR for bacterial microbiome

The abundance of the Eubacteria domain in seed and maize root was measured by real-time PCR from the 16S rRNA. For this, the total DNA of the samples was extracted according to the methodology mentioned in the previous topic (CTAB method) and amplified from primers 926F (AAACTCAAAKGAATTGACGG) and 1062R (CTCACRRCACGAGCTGAC) (De Gregoris et al., 2011). PCR was performed in triplicate and with 15 μL of a reaction containing DNA (100 ng for seed and 40 ng for root), 7.5 μL of SYBR Green (Promega), 0.5 μL of each primer (10 μM) and water. The reaction conditions were 5 min incubation at 95 ° C, 40 cycles of 15 s at 95 ° C and 1 min at 60 ° C in Step-One-Plus Real-Time PCR System (Applied Biosystems). The standard curve was generated by diluting the DNA of the bacterium *Escherichia coli* ATCC 25922 in series of 10^2^ – 10^−8^ (20 – 2 × 10^9^ ng of DNA). *E. coli* was grown in NB liquid medium (180 rpm, at 30 ° C) and had its DNA extracted with Wizard Genomic DNA Purification Kit (Promega). The number of bacteria in the seed and root was calculated based on the values of Ct (cycle threshold) and the standard curve (Staroscik, 2004).

### Mobilization of seed-maize reserves during germination

To measure the mobilization of reserves during the germination of maize, seeds were treated and germinated according to item 2.1, with some modifications, including the soaking of NDS in sterile distilled water for 35 min. At the same time, DS was immersed for an equal period in alcohol/hypochlorite. Biochemical analyzes on maize were carried out in three distinct stages: 1°) Imbibition (seeds sampled after disinfection); 2°) End of germination (embryonic axis collected 24 (for SND) and 48 h (for SD) after radicle emission (5 mm); 3°) Seedling stage (root collected after 5 days of germination). In stage 2, the embryonic axis was collected at different times because disinfection reduces the speed of seed germination (Fig 2). The samples collected in the 3 stages were macerated in liquid nitrogen and analyzed, in triplicate, for protein (Smith et al., 1985), reducing sugar (Miller, 1959), glucose, triglycerides and alpha-amylase activity (Bioclin® K082, K117, K003), according to the cited protocols. The results were analyzed by ANOVA, followed by Tukey test (p ≤ 0.05).

### Effect of bacterial microbiota on biocontrol

To evaluate the potential of maize seed-borne bacteria in the control of phytopathogenic fungi, disinfected, and non-disinfected seeds were assayed as described in the 2.1. follow inoculation with *Penicillium* sp. This fungus was initially isolated from another maize variety (*Z. mays* var. SHS5050) after inhibiting 100% of its germination and affected seed reserve remobilization in the previous study with the same experimental setup under the axenic condition described herein.

Once isolated, *Penicillium* sp. was grown in solid potato-dextrose-agar (PDA) in BOD (30 °C; 7 days), followed by new growth in NB liquid medium (180 rpm, at 30 °C) and inoculation in NDS and DS (150 μL per seed; OD595 = ~1; 10 seeds per plate; 4 repetitions). After five days, germination rate and seedling growth (length and mass) were evaluated, while scanning microscopy was used to characterize fungal colonization at the root of the different treatments (methodologies described in topics 2.1 and 2.3, respectively). The results were submitted to analysis of variance (ANOVA) and the means compared by the Tukey test (p ≤ 0.05).

## Results

To assess the role of seed-borne bacteria in maize, a time-course chemical seed-disinfection assay with sodium hypochlorite was carried out to remove most of the microbial community without impairing the germination process. Thus, we selected the time of 30 min of seed immersion in 1.25% sodium hypochlorite solution. Disinfecting the seeds for 30 min removed most of their bacteria (Fig. 1A), without affecting the germination percentage of the maize (Fig. 2C). Live/Dead® viability tests confirmed that the disinfection reduced the number of viable bacteria cells compared to the non-disinfected treatment (Fig. 1B). Both treatments showed green fluorescence (live cells), with few visible (red) dead cells (Fig. 1B).

**Fig. 1.**
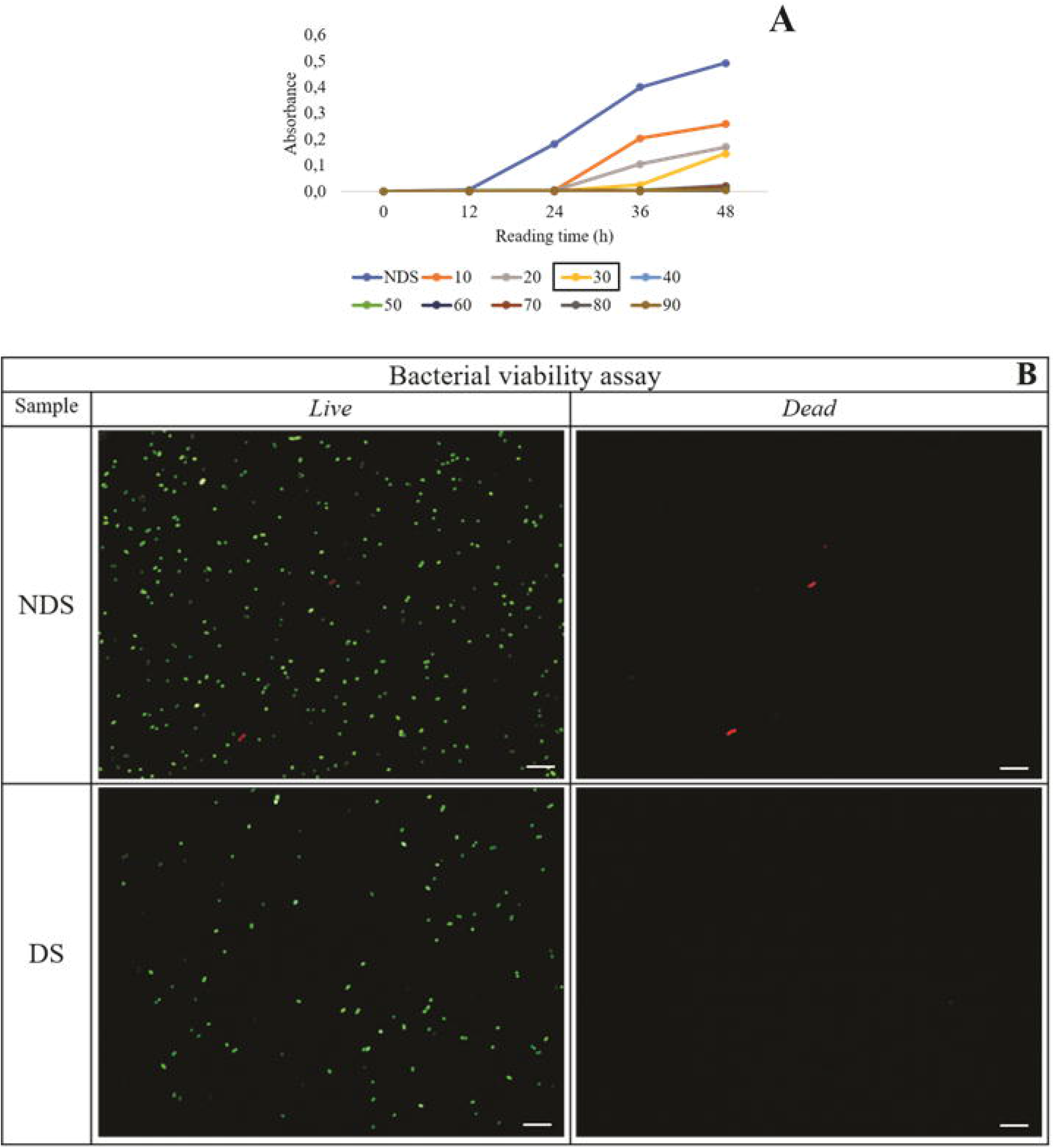
Bacterial growth reported as optical density increase in NB liquid medium with maize seeds submitted to different disinfection times in sodium hypochlorite (1.25% for 10, 20, 30, 40, 50, 60, 70, 80 and 90 min) (A) and fluorescence live/dead bacterial viability assay images of seeds non-disinfected (NDS) and disinfected (DS) (B). Scale bar: 10 μm.

**Fig. 2.**
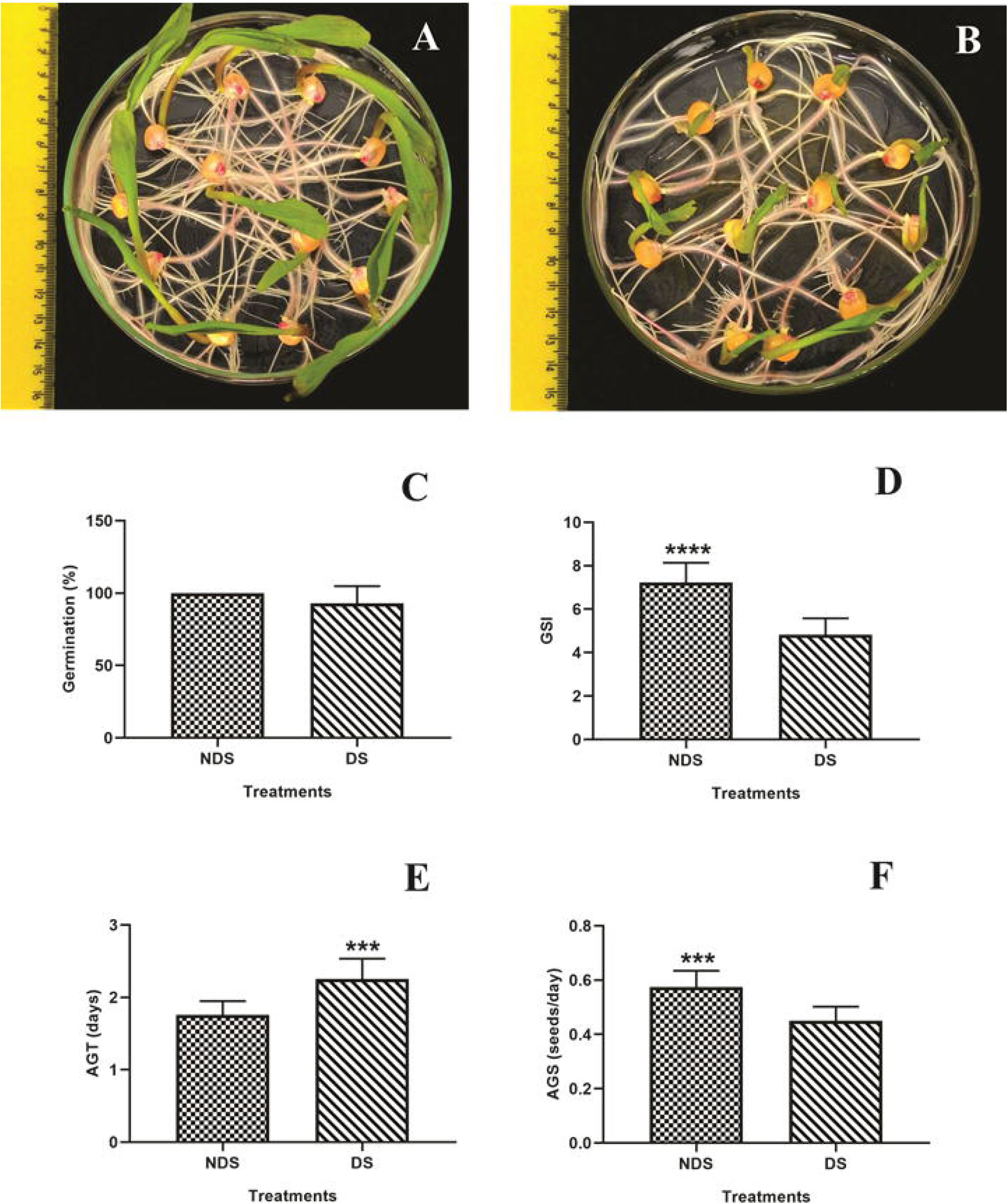
Germination of non-disinfected (NDS; A) and disinfected (DS; B) maize seeds. Germination percentage (C), germination speed index (D), average germination time (E), and average germination speed (F). *Significant difference between treatments according to the Tukey test (p ≤ 0.05).

The seed-disinfection did not affect the germination percentage of the maize (Fig. 2C) but reduced the germination speed (Fig. 2D and 2F) and increased the time necessary for the seed to germinate (Fig 2E).

Seed-disinfecting treatment significantly reduced diazotrophic bacteria population associated to emerged radicle of maize seedling (5 days after emergence) in LGI semi-solid N-free medium (sucrose as a C-source) and dramatically reduced the diazotrophic bacteria population grown in JNFb semi-soild N-free medium (malic acid as C-source) to no detected level (Fig. 03). For total bacteria in NB solid medium, it was shown no significant decrease in the root seedling population in disinfect seeds. The same trend was obtained by population size of root segments overnight enriched in NB liquid medium (Fig 03)

**Fig. 3.**
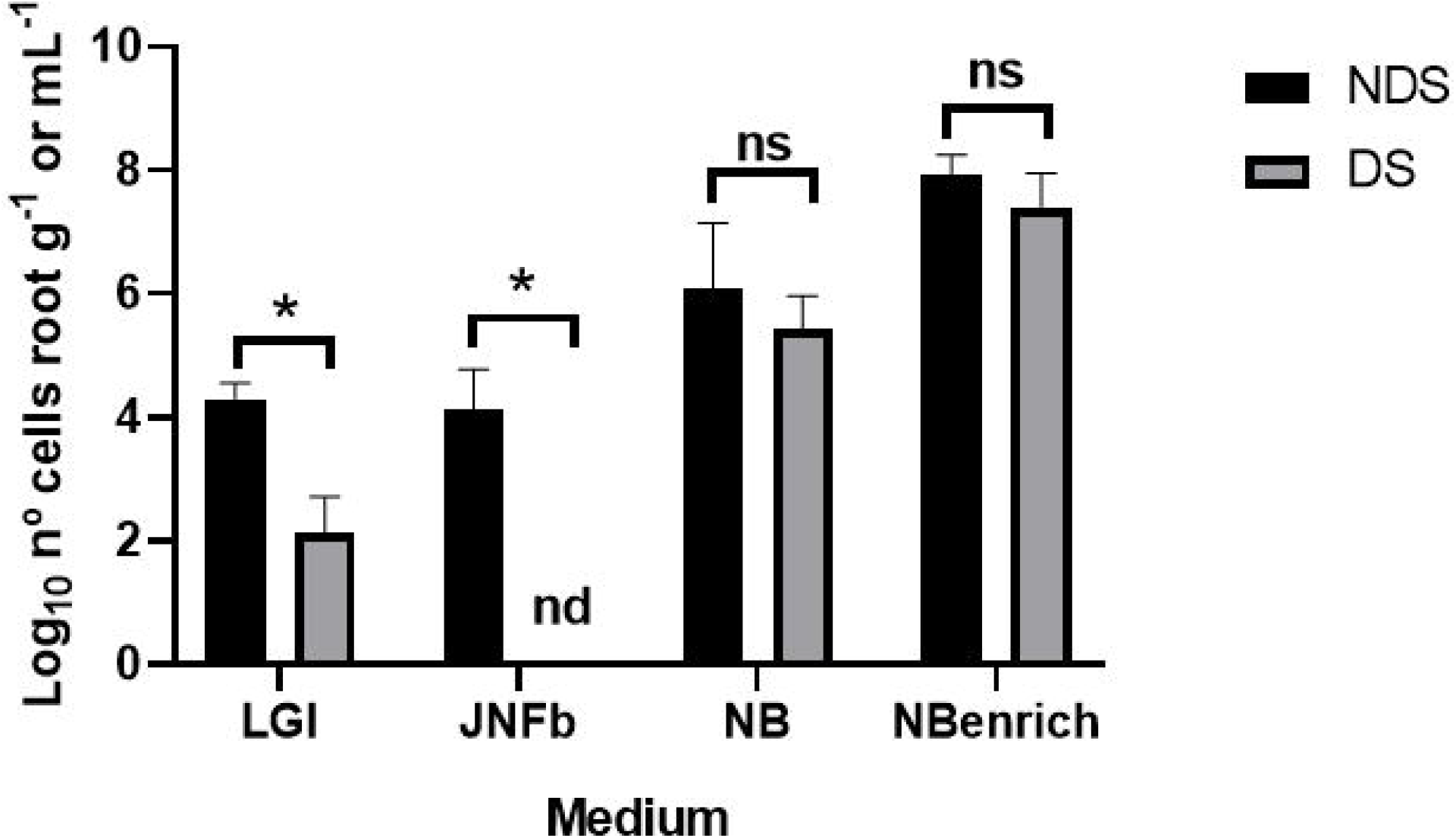
Influence of seed disinfection (DS versus NDS) on bacterial population counts associated to five days emerged radicle of maize seedling (*Z. mays* DKB 177) under axenic conditions recovered in different culture media. Maize seeds non-disinfected (NDS) and disinfected (DS). nd: not detected. Positive growth in LGI (sucrose as C-source) and JNFb (malic acid as C-source) semi-solid N-free media means the formation of a white sub-superficial white pellicle Data for diazotrophic expressed as log_10_ n° cells. g^−1^ root with three biological replicates. Data for total bacteria expressed as log_10_ n° CFU. g^−1^ root or mL^−1^ in NB solid medium.

Elongation/differentiation zone cross-section of the emerged radicle from non-disinfected (Supplementary Fig 1A1-A3) and disinfected maize seeds (Supplementary Fig. 1B1-B3) were viewed under light (LM) and transmission electron microscopy (TEM). The tissue system organization of the root tissue can be visualized in images of LM (Supplementary Fig. A1). Under TEM, the regular orientation of the plant cell wall, cytoplasmatic organelles, and the presence of prominent vacuoles in both treatments were noticed (Supplementary Fig. A1). Thus, it seems unlikely that the hypochlorite affected the root anatomy organization of maize root seedling.

Scanning electron microscopy (SEM) was used to characterize the niche occupancy by the bacterial community in water-imbibed seeds before germination (Fig. 4). It was not possible to distinguish population density differences of bacteria in the pericarp coat (Fig. 4A1 and 4B1) and endosperm region (Fig. 4A2 and 4B2) from disinfected and non-disinfected seeds. After 48 h-germination, with radicle protrusion that emerged throughout the micropyle, it was noticed a remarkable densely bacteria populations at the bottom of the radicle surface in both treatments (Fig. 4A3-A4 and 4B3-B4).

**Fig. 4.**
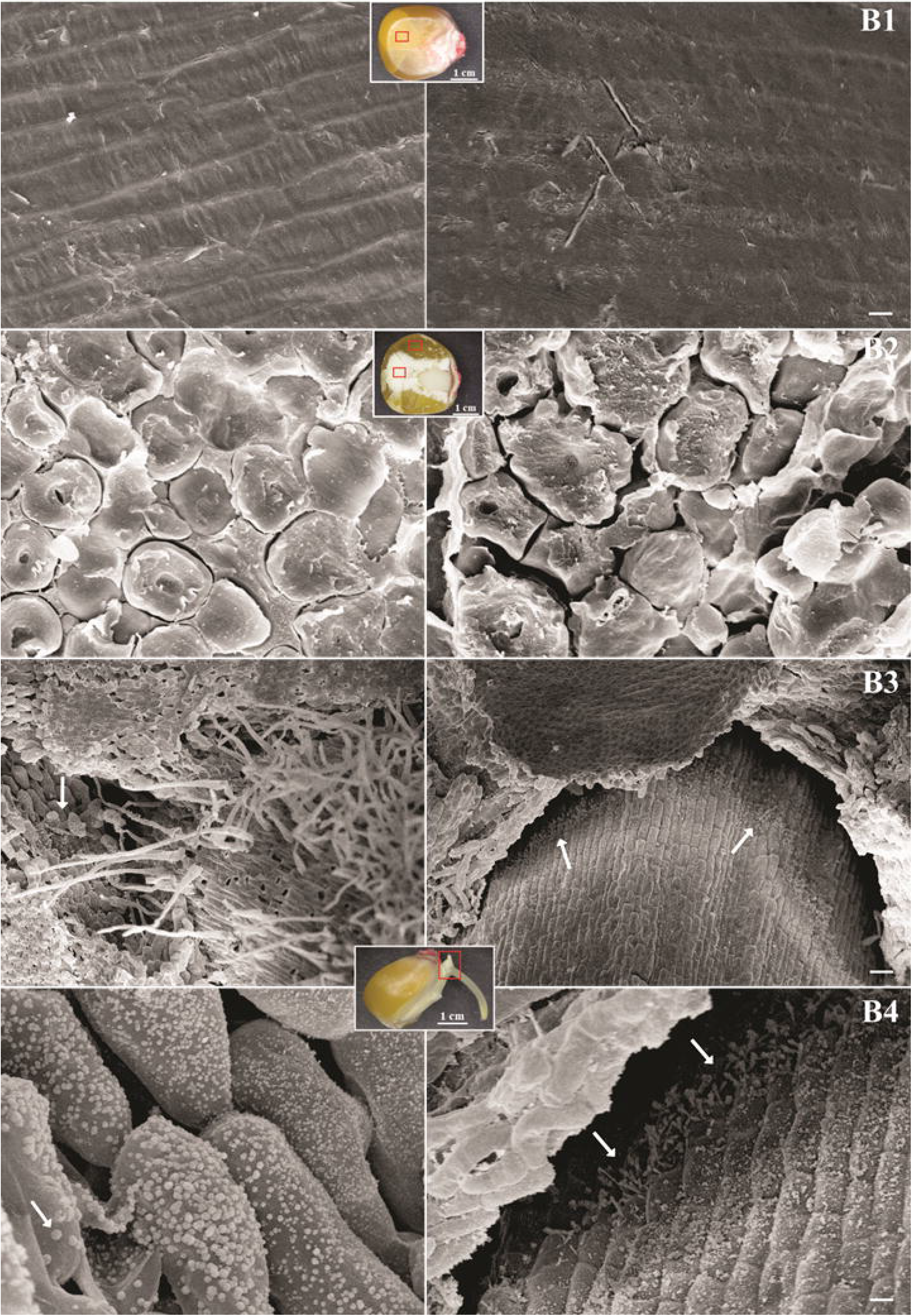
Colonization of maize seeds by its microbiota. Bacterial cells were visualized by scanning electron microscopy (SEM). Maize seeds non-disinfected (A) and disinfected (B). Seed regions: pericarp (A1 and B1), endosperm (A2 and B2), and radicle (A3, A4, B3, and B4). Biofilms and small bacterial aggregates are indicated by white arrows. Bars represent the following scales: panel B2: 3 μm; A2: 10 μm; A1, A4, B1, and B4: 20 μm; A3: 100 μm; B3: 200 μm.

After 5 d-germination, bacterial cells were viewed under LM and SEM. In root-segments stained with TTC, few bacteria aggregates were seen in the lateral root emission region (Fig 5A1 and 5B1). In contrast, no bacteria were visualized directly in the root tip surface of the disinfected and non-disinfected treatments (Fig 5A2 and 5B2). SEM observations showed many single to small aggregate bacterial cells distributed over root hair zone of NDS-treated (Fig. 5A3), while DS bacteria were viewed as larger aggregate with less frequent colonization pattern over root hair zone (Fig 5B3). At the elongation zone of DS, bacteria cells were main visualized as single cells, while aggregated or biofilms of bacteria community were observed in the NDS (Fig 5A4 and 5B4). In the root tip segment, the pattern colonization was similar for both (NDS and DS) with bacteria community more frequently organized as small aggregates (Fig 5A5 and 5B5).

**Fig. 5.**
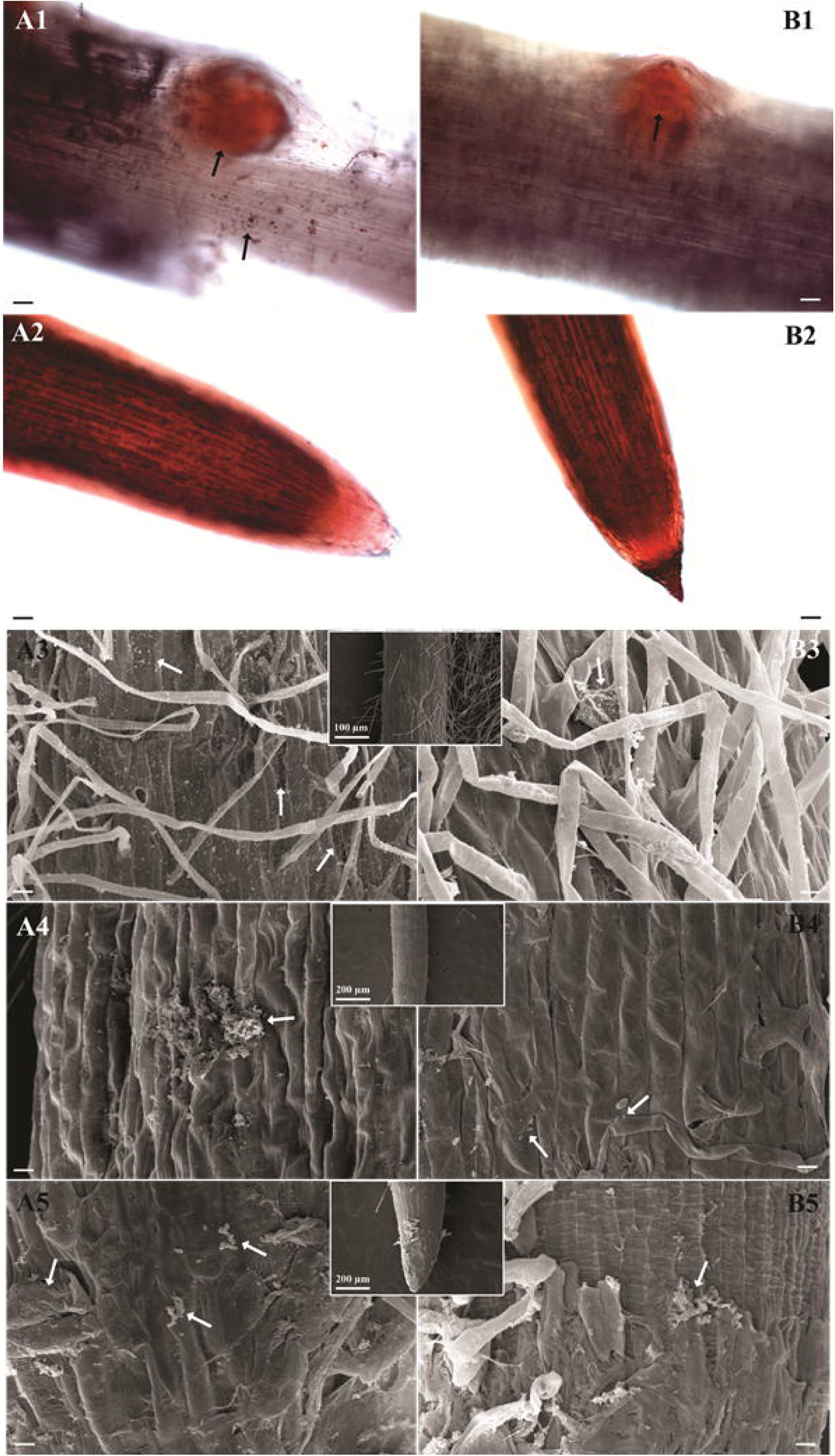
Bacterial colonization of maize root visualized by light (LM; root stained with TTC) and scanning electron microscopy (SEM). Maize seeds non-disinfected (A) and disinfected (B). Root regions: mitotic sites (A1 and B1), root hair (A3 and B3), the zone of elongation (A4 and B4), root cap (A2, A5, B2, and B5). Bacteria are indicated by arrows. Bars represent the following scales: panel A1, A2, B1, and B2: 10 μm; A3-A5 and B3-B5: 20 μm.

By sequencing and analyzing the16S rRNA, we characterize the bacteriome of the transition phase of the seed to root seedling emergence under axenic conditions. Data sequencing of 5-h embedded seed resulted in 587 reads in total, with an average sample coverage of 53% (Supplementary Table 1). PCoA analysis for seed bacteriome showed distinct clustering between some samples independent of disinfected and non-disinfected seeds (Fig. 6A). The seed bacteriome showed similar diversity (p = 0.66) (Fig. 6B) and did not differ in structure (p = 0.6; R^2^ = 0.195) (Supplementary Table 2) after the disinfection process.

**Fig. 6.**
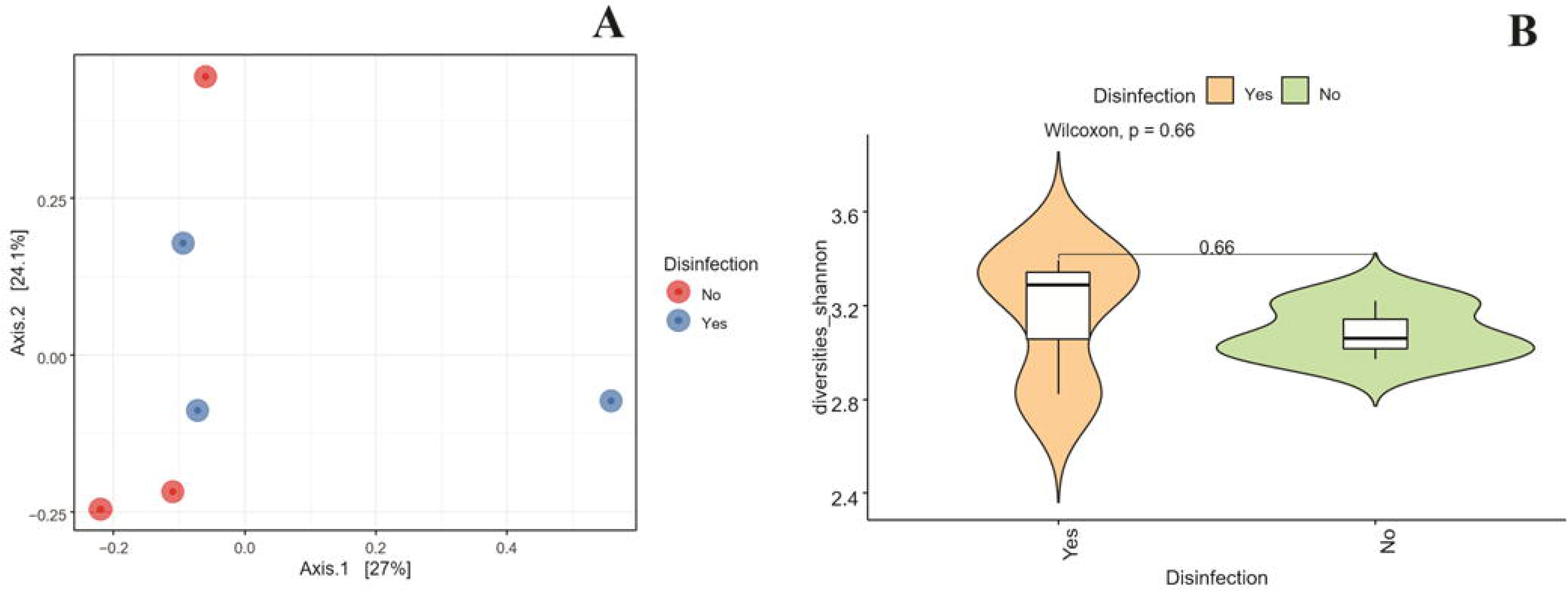
Principal coordinate analysis (PCoA) plot and alpha diversity of the bacteriome associated with non-disinfected (No) and disinfected (Yes) seeds in axenic systems.

Venn diagram revealed a low number of OTUs in non-germinated embedded seeds, with two unique OTUs for non-disinfected seeds and 3 for disinfected seeds (Fig. 7A). Only 2 OTUs were shared between treatments (Fig. 7A). In the bacteriome associated with the seed, we identified ten genera from 3 different phyla, attributed to Proteobacteria (7 genera), Firmicutes (2 genera), and Actinobacteria (1 genus) (Fig. 7B). Among the ten genera, *Azospirillum*, *Acinetobacter,* and f_Enterobacteriaceae_922761 (unassigned genus) were more abundant in non-disinfected seeds, while *Acinetobacter* and *Staphylococcus* were abundant in seeds treated with hypochlorite. Other genera were detected in low numbers (Fig. 7B). Disinfection appears to reduce the relative abundance of f_Enterobacteriaceae_922761, *Azospirillum,* and *Acinetobacter* in the seed. However, the analysis of differential abundance revealed that no taxon was removed by the hypochlorite (Fig. 7B).

**Fig. 7.**
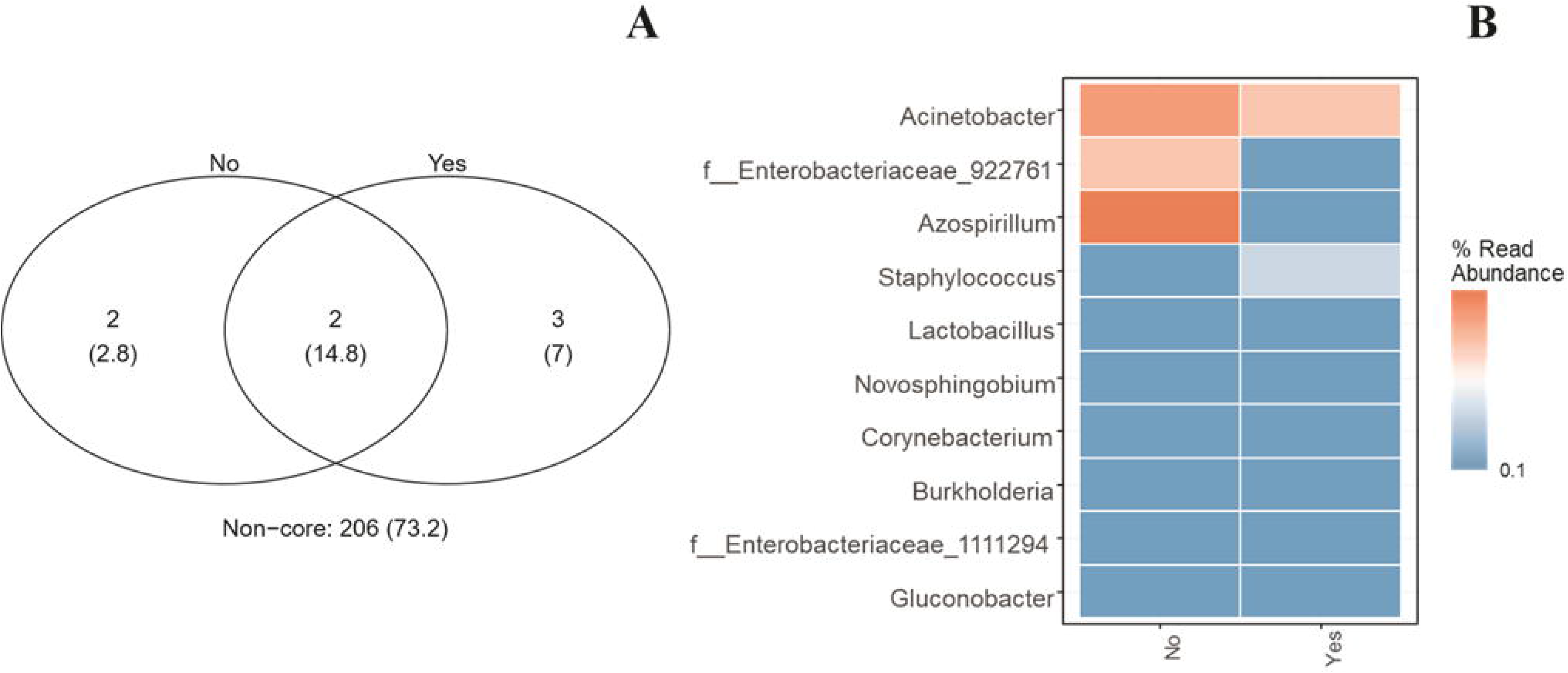
Venn diagram with shared OTUs and heat map with the taxonomy of the most abundant genera of non-disinfected (No) and disinfected (Yes) seeds maintained in axenic conditions.

Five days after seed germination, the sequencing of the maize root bacteriome resulted in 2,22923 reads in total and average sample coverage of 86% (Supplementary Table 3). PCoA analysis showed that the beta dispersion of the samples did not differ between the emerged roots of NDS and DS (Fig. 8A), while Permanova indicated that disinfection was not significant for grouping biological replicates (p = 0.384; R2 = 0.091) (Supplementary Table 4). The bacterial diversity of the root was also not altered by disinfection (p = 1.0) (Fig. 8B). We observed a high overlap of OTUs between bacteria communities associated with emerged roots of NDS and DS, with 10 OTUS in common (Fig. 9A). Only 3 and 6 exclusive OTUS were observed for NDS and DS, respectively (Fig. 9A). In the root bacteriome, we identified ten genera, nine classified in the phylum Proteobacteria and 1 in the phylum Bacteroidetes (Fig. 9B). Genus differences include a greater abundance of *Pseudomonas*, *Acinetobacter*, *Achromobacter*, *Methylobacterium,* and *Novosphingobium* in the roots of disinfested treatment (Fig. 9B). Interestingly, roots from the disinfected treatment were densely colonized by bacteria abundant in NDS roots, plus the genera *Zoogloea*, f_Comamonadaceae_942852 (unassigned genus), *Ralstonia*, *Sediminibacterium* and f_Oxalobacteraceae_1033018 (unassigned genus) (Fig 9B). We found 3 OTUs that differed in abundance in disinfected versus non-disinfected seeds and were attributed to the genera *Acinetobacter* and *Ochrobacterium* (Supplementary Table 5). Within *Acinetobacter*, only the species *A. rhizosphaerae* was identified.

**Fig. 8.**
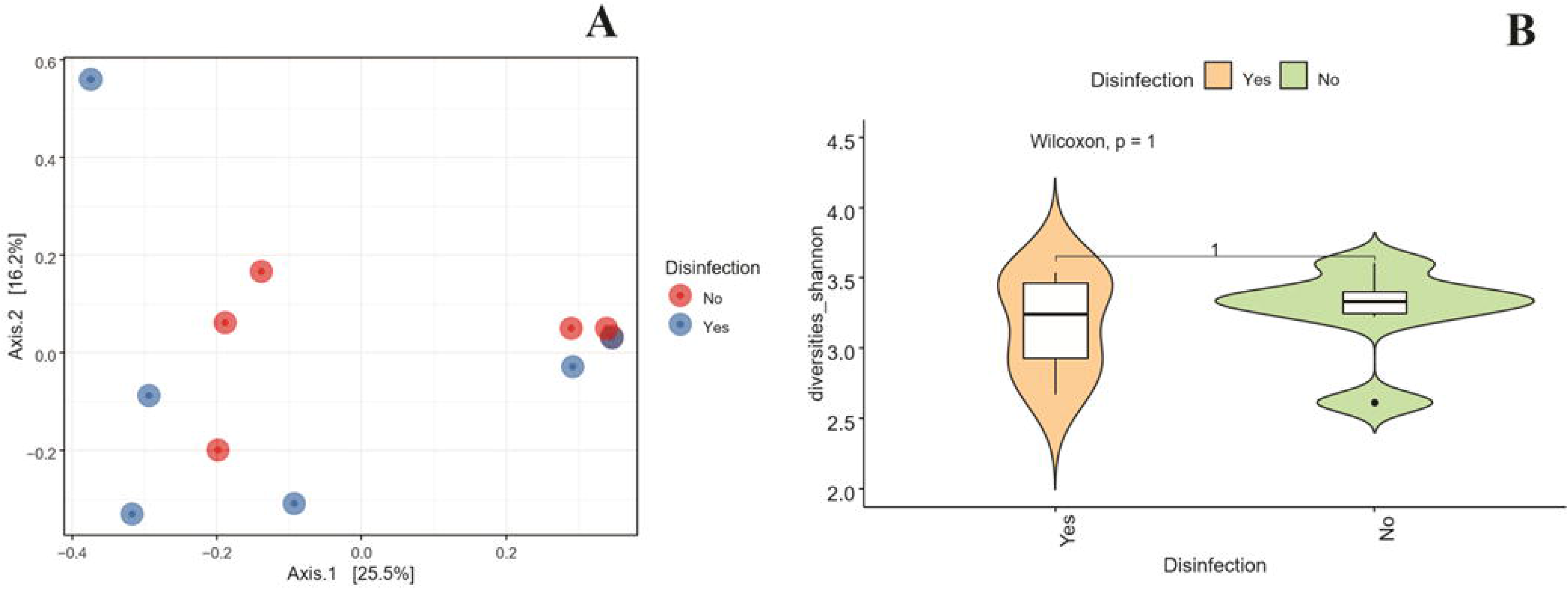
Principal coordinate analysis (PCoA) plot and alpha diversity of the bacteriome associated with the root of non-disinfected (No) and disinfected (Yes) seeds in axenic systems.

**Fig. 9.**
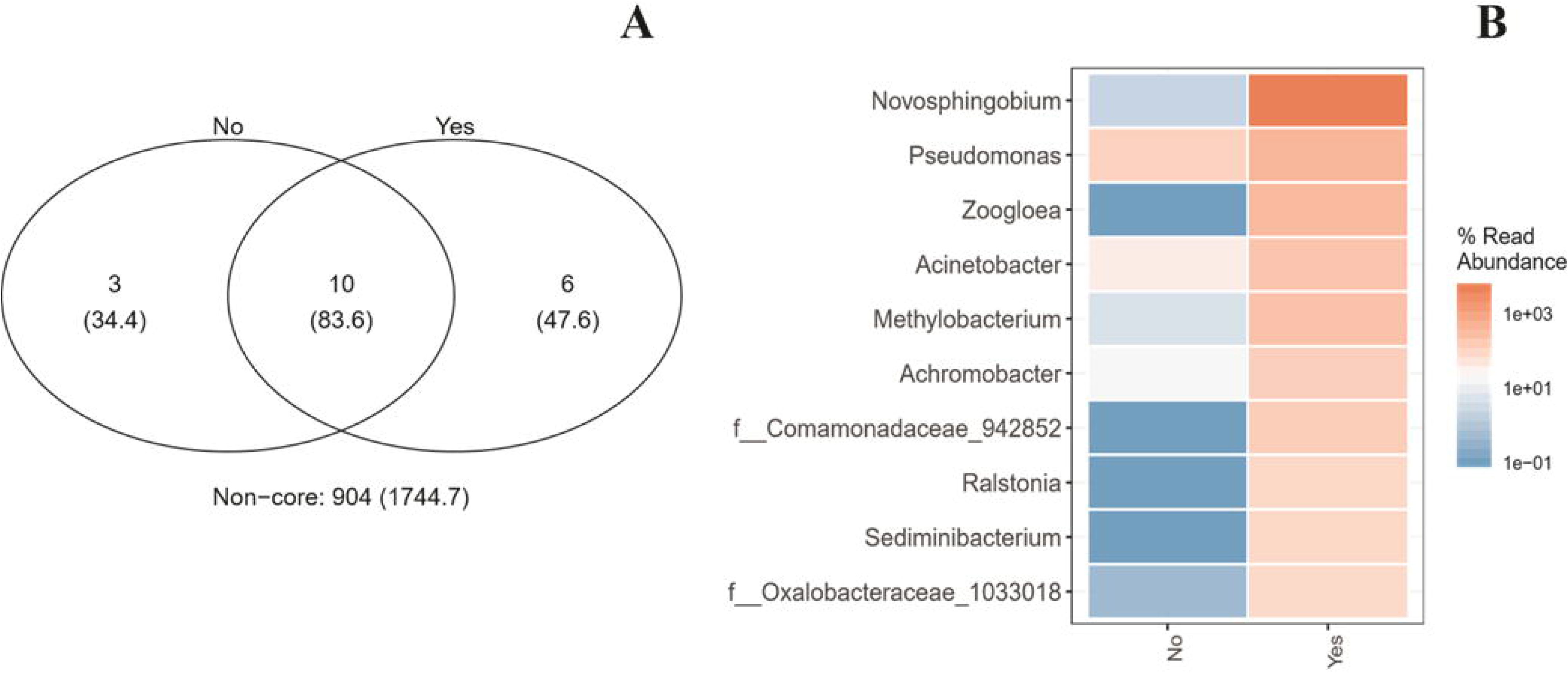
Venn diagram with shared OTUs and heat map with the taxonomy of the most abundant genera in roots of non-disinfected (No) and disinfected (Yes) seeds maintained in axenic conditions.

The quantification of bacteria via real-time PCR identified a similar number of cells/ng of DNA in seeds (Fig. 10A; NDS: 7,356 log cell; DS: 7,287; p ≥ 0.05), and roots (Fig. 10B; NDS: 5,360 log cell; DS: 5,208; p ≥ 0.05) disinfected and not disinfected.

**Fig. 10.**
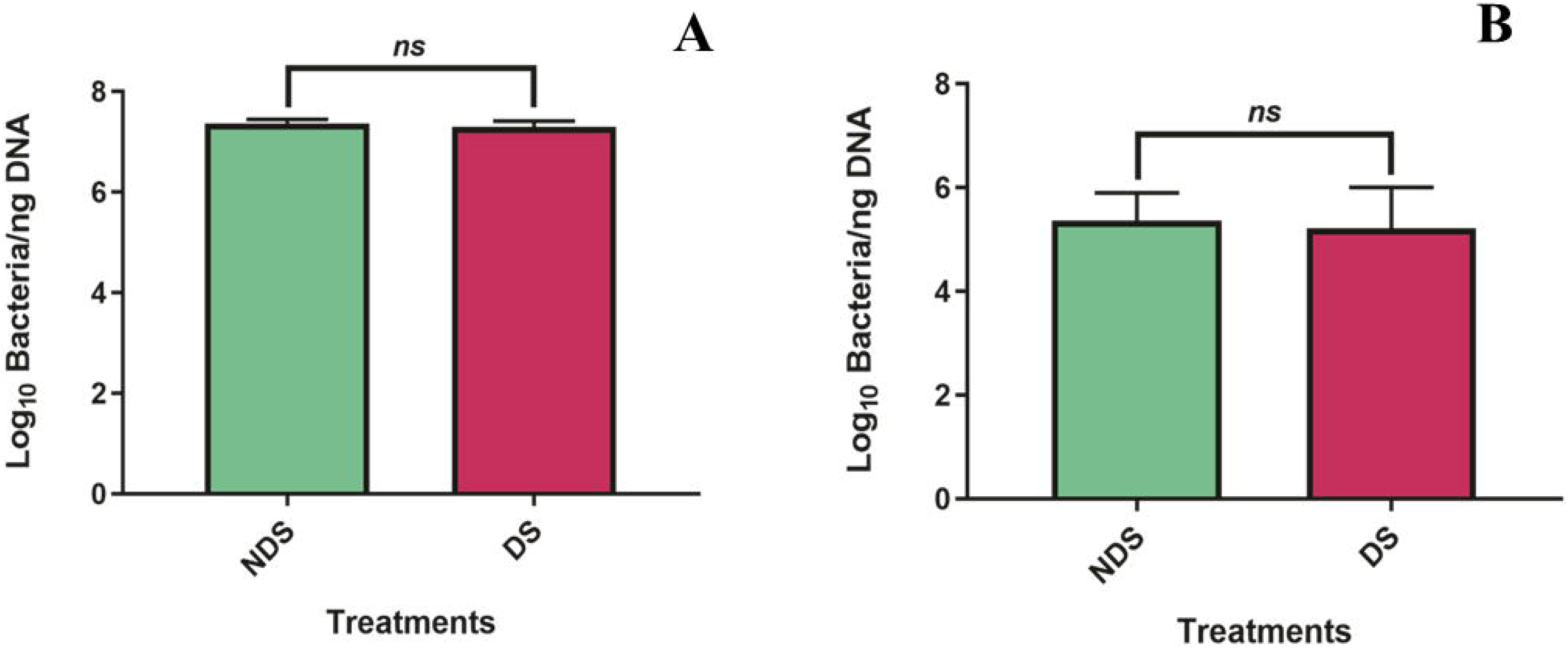
Quantification of bacteriome by real-time PCR in non-disinfected (NDS) and disinfected (DS) roots (A) and seeds (B). *Significant difference between treatments according to the Tukey test (p ≤ 0.05).

Significant differences in the mobilization of maize reserves were only observed in the degradation of triglycerides of the embryonic axis (Fig. 11C) and activity of the alpha-amylase enzyme to the root (Fig. 11E). In both cases, the results were superior in the non-disinfected treatment. The protein, glucose, and reducing sugar content did not differ between treatments (NDS and DS) of the seed compartment, embryonic axis, and root (Fig. 11A, 11B, and 11D). In general, the mobilization of reserves changed as germination progressed, with significant differences between the analyzed stages (Supplementary Table 6).

**Fig. 11.**
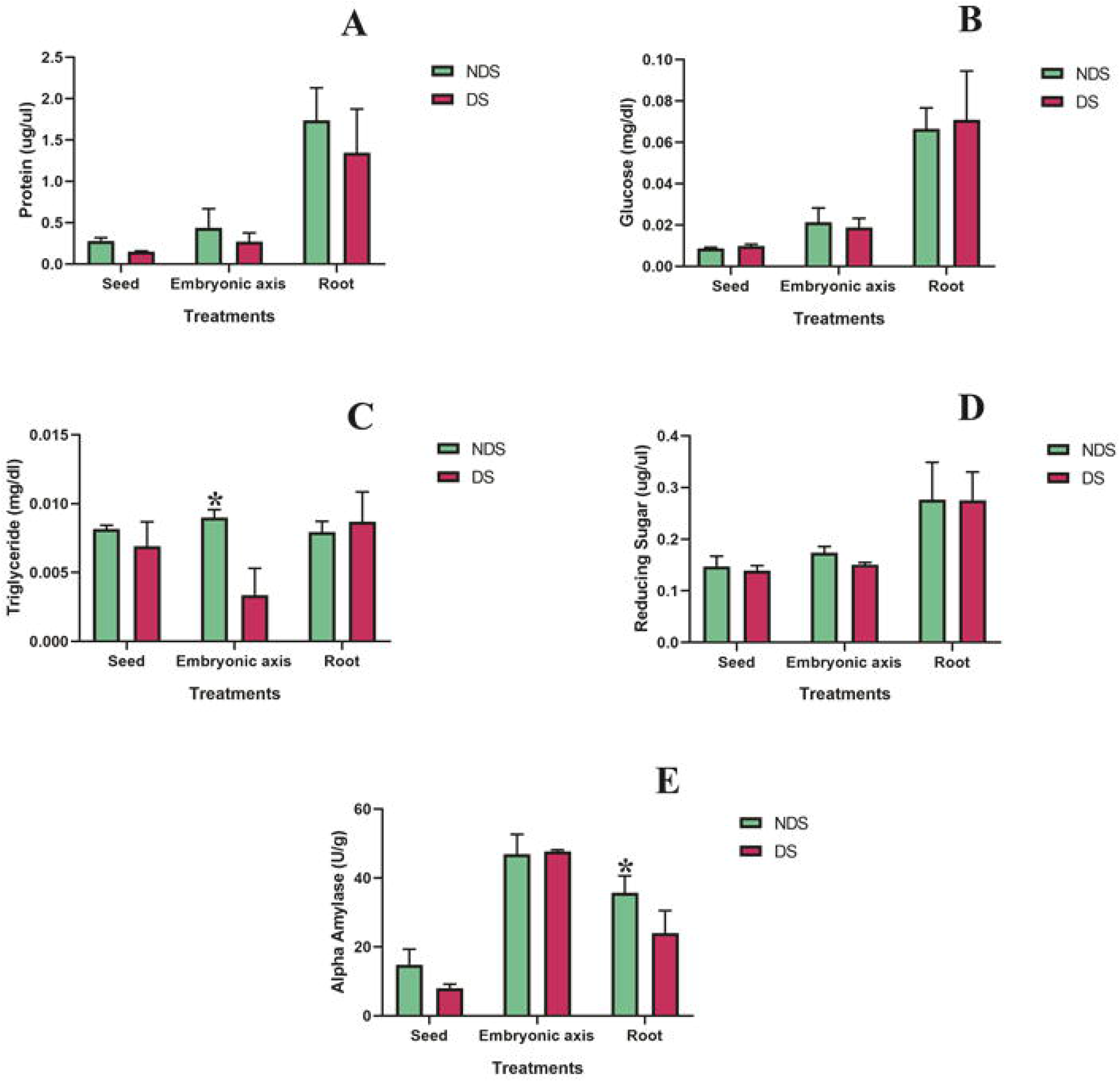
Dosage of protein (A), glucose (B), triglyceride (C), reducing sugar (D), and alpha-amylase activity (E) in seed, embryonic axis, and root non-disinfected (NDS) and disinfected (DS). *Significant difference between treatments NDS and DS (within each stage) according to the Tukey test (p ≤ 0.05).

We also evaluated if the observed changes in the seed-borne bacteria community interfere with tolerance against seed-borne phytopathogenic fungi. Maize germination was tested after seed disinfection and inoculation of *Penicillium* sp. In Fig. 12, we observed that fungus significantly reduced the percentage (C) and the germination speed (D) of the disinfected seeds, which had part of their microbiota re-shaped by the action of the hypochlorite. Due to the action of the fungus, many disinfected seeds decay before or just after germination (Fig. 12B2). When not disinfected, the seeds that received the fungus germinated normally (Fig. 12C and 12D), with percentage and speed equal to the control (without inoculation of the fungus). The average germination speed and time did not differ between treatments (Fig. 12E and 12F).

**Fig. 12.**
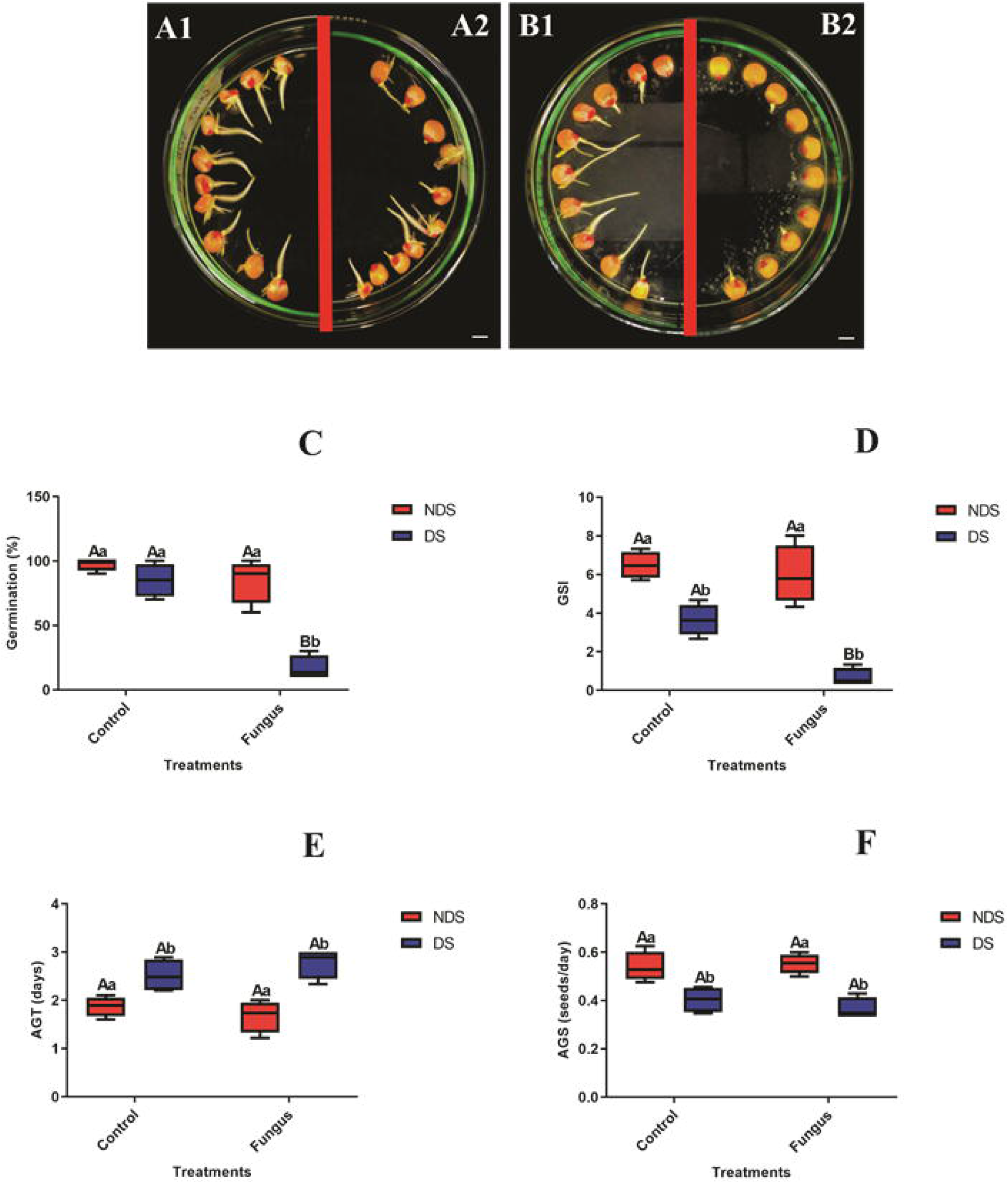
Germination of non-disinfected (NDS; A1 and B1) and disinfected (DS; A2 and B2) maize seeds inoculated (B1-B2) or not (A1-A2) with fungus *Penicillium* sp. Germination percentage (C), germination speed index (D), average germination time (E), and average germination speed (F). Different capital letters indicate significant differences for the inoculation factor (Control x Fungus), and lowercase letters indicate significant differences for the disinfection factor (NDS x DS) according to the Tukey test (p ≤ 0.05).

The growth of germinated seedlings from disinfected seeds and inoculated with *Penicillium* sp was drastically reduced, affecting the length of the aerial part (LAP) and root (LR), as well as the fresh root mass (FMR) (Fig. 13C, 13D, and 13F). These results were significantly inferior to the non-disinfected seeds challenged with the phytopathogenic fungus and the disinfected-without-inoculation control (Fig. 13). It is noteworthy that disinfected-inoculated seedlings rotted, showing brown spots over the root axis (Fig 13B2). Seedlings of seeds that were not disinfected and inoculated with fungus grew normally for all characteristics analyzed (except FMAP), with LAP, LR, and FMR values close to the control without inoculation (Fig 13C, 13D, and 13F). Only the fresh weight of the aerial part (FMAP) did not differ between treatments (13E).

**Fig. 13.**
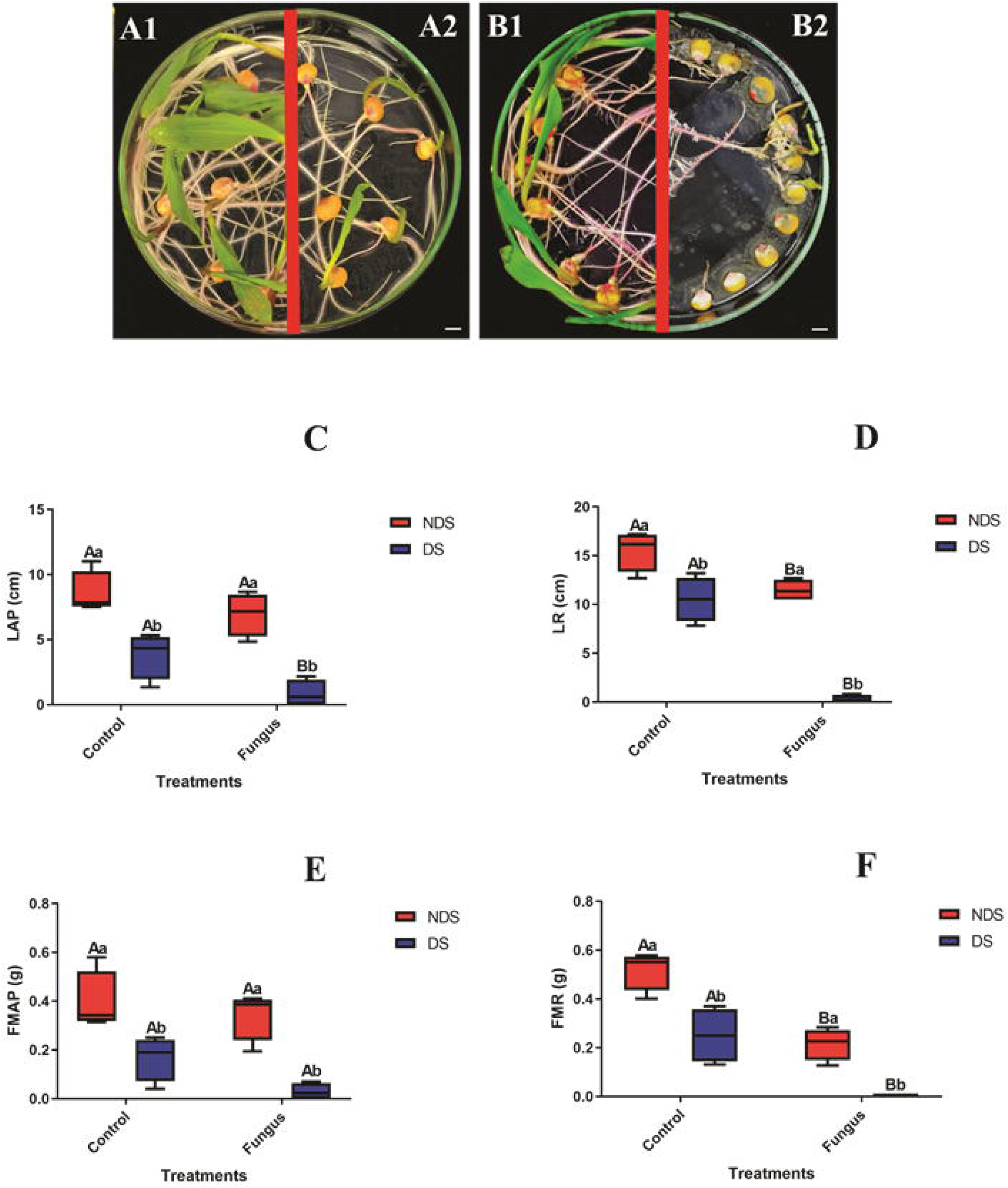
Growth of maize germinated of non-disinfected (NDS; A1 and B1) and disinfected (DS; A2 and B2) maize seeds inoculated (B1-B2) or not (A1-A2) with fungus *Penicillium* sp. Shoot length (C; LAP) and root (D; LR), shoot fresh weight (E; FMAP), and root (F; FMR). Different capital letters indicate significant differences for the inoculation factor (Control x Fungus), and lowercase letters indicate significant differences for the disinfection factor (NDS x DS) according to the Tukey test (p ≤ 0.05).

Scanning electron microscopy (SEM) identified the presence of fungal hyphae in maize roots infected with *Penicillium* sp (Fig. 14). Mycelia densely colonized the elongation/differentiation zone (Fig. 14B3 and 14B4) and the root tip region(Fig. 14B5) of the germinated roots of the disinfected seeds. The lesser density of bacteria and yeast aggregates were observed in the infected region (Fig. 14B4). In the roots of non-disinfected seeds, SEM showed few hyphae in the elongation/differentiation zone (Fig. 14A3 and 14A4), and no hyphae were viewed in the root tip region (Fig 14A5). In this treatment, it was observed a high number of bacteria attached to the root surface in monolayer, interacting with fungus hyphae (Fig. 14A4). *Penicillium* sp., yeasts, and bacteria colonized root areas with lateral root emergence, in the disinfected treatment (Fig. 14B1 and 14B2), while bacterial biofilms were seen in the non-disinfected treatment in the same niche (Fig. 14A1 and 14A2).

**Fig. 14.**
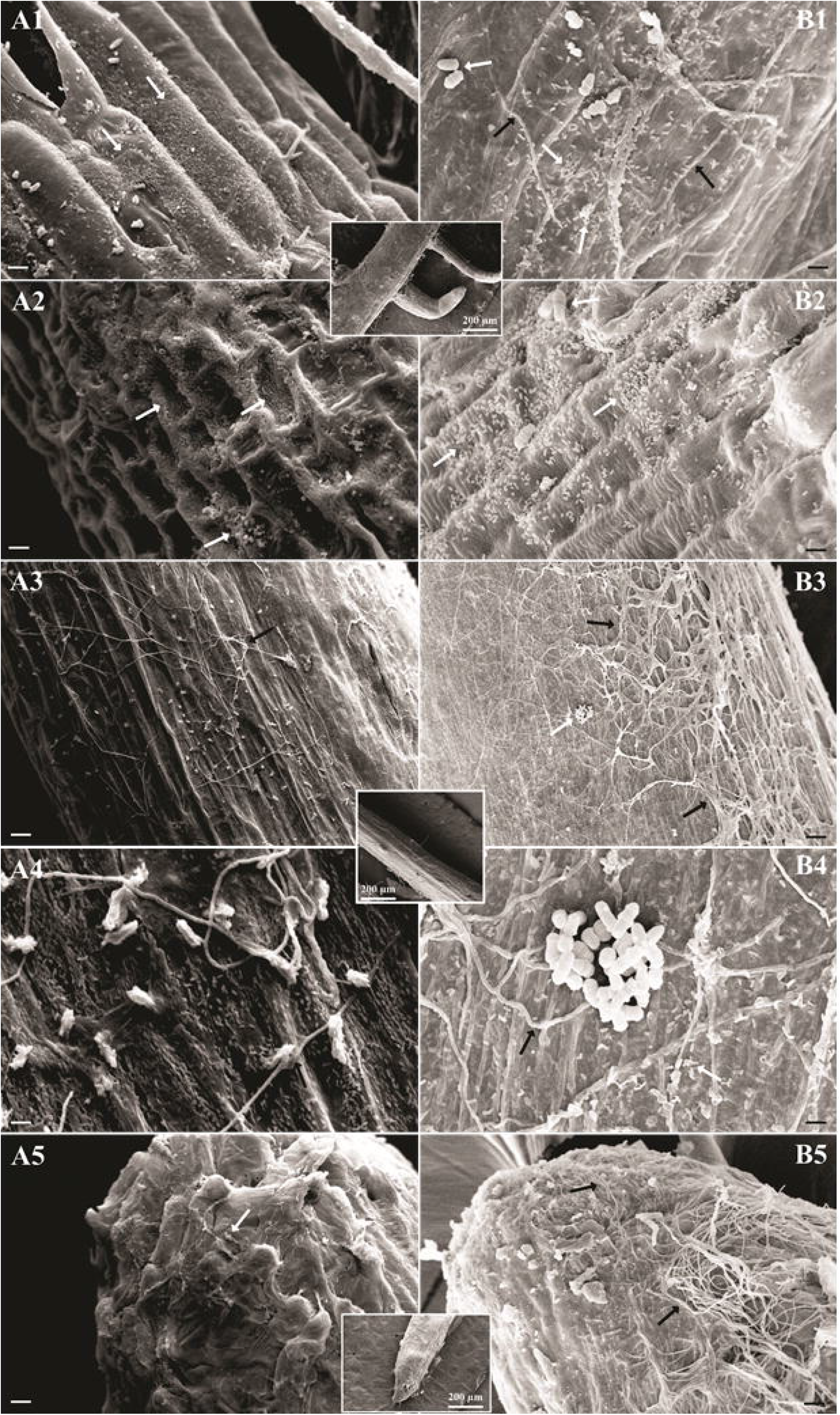
Colonization of maize root by *Penicillium* sp. Fungi were visualized by scanning electron microscopy (SEM). Maize seeds non-disinfected (A) and disinfected (B). Root regions: lateral root (A1, A2, B1, and B2), the zone of elongation (A3, A4, B3, and B4), root cap (A5 and B5). Bacteria/yeasts and filamentous fungi are indicated by white and black arrows, respectively. Bars represent the following scales: panel A2, A4, B1, B2, and B4: 10 μm; A1, A3, A5, B3, and B5: 20 μm.

No filamentous fungal tissue was observed in uninoculated seedling roots (Supplementary Figure A2). In control, bacteria and yeasts were seen colonizing the root tissue in isolation or small aggregates (Suppl. Fig. A2). A higher density of bacteria was detected in the root hood of the non-disinfected treatment (Suppl. Fig. A2).

## Discussion

For a long time, “sterile” seeds were considered healthy, which contributed to the development of disinfection methods (chemical, mechanical, physical, and biological) in order to remove their “pathogens” (Berg and Raaijmakers, 2018). However, in recent years, studies based on “omics” have shown that the seeds harbor diverse and mostly beneficial microbial communities (Berg and Raaijmakers, 2018). In the present work, we confirmed that axenically germinated maize seeds host several bacteria taxon, which was located, quantified and identified by microscopic analysis, real-time PCR, counting in the culture medium, and sequencing. Also, it was demonstrated that from seed to seedling transition, there was a substantial increase in size and complex of the bacteria community structure, whose functionalities remain to be elucidated.

In this axenic study, soaking maize seeds in sodium hypochlorite solution (1.25%, 30 min) proved its antimicrobial effect through the Live-Dead viability assay and population estimation by count in the culture medium. However, microscopy analysis of the water embedded seed and emerged roots from germinated maize seeds revealed that the chemical disinfection reduces but does not remove all bacteria from the seed. Pieces of evidence for this are images of SEM and LM with no visible differences in the bacterial density of the treatments (NDS *versus* DS).

After reducing the number of maize bacteria with disinfection, a delay in seed germination speed and seedling growth was observed that reinforces the idea that some bacteria borne in seeds are essential for these physiological processes. Other studies have shown that chemical disinfection (Irizarry and White 2017; Verma et al., 2017; Verma et al., 2018; Verma and White, 2018) and thermal treatments (Holland, 2016; Holland, 2019) have slowed germination and growth of rice, soybean, beans, and millet. It is worth mentioning that disinfection did not affect the germination percentage of maize and did not structurally alter the plant cell and tissue viewed by transmission electron microscopy. Therefore, it seems unlikely that delays in the speed of germination and growth are attributed to sodium hypochlorite.

The idea that disinfection can reduce, but not remove all bacteria from maize, was confirmed by sequencing the 16S rRNA, since there were no significant differences between treatments (NDS versus DS) for Permanova and Shannon diversity, besides the significant number of shared OTUs. Differences in sequencing were restricted to the distinct taxonomic composition between the seed and root compartments, which shared only two genera (*Acinetobacter* and *Novosphingobium*). Within the studied compartments, the variation between treatments is only quantitative; that is, it is based on the abundance of taxa, not on their presence or absence, which justifies the fact that only 3 OTUs have been removed by disinfection (*Acinetobacter* (2) and *Ochrobacterium* (1)). In the seed, the reduction of the genera f_Enterobacteriaceae_922761 (unassigned genus), *Azospirillum,* and *Acinetobacter* after disinfection can be related with later emergence of several new taxa (9 genera in all) activated at the emerged root during germination, with emphasis on the genus *Novosphingobium*. These findings suggest that the reduction of dominant genera in the seed reduced the competition for niches or resources in the root, allowing colonization by other bacterial groups (Hardoim, 2019). However, these new groups do not contemplate or contemplate a reduced number of bacteria that promote germination and growth, which affected the development of maize.

Once the bacteriome of maize is characterized, it remains to elucidate how these microorganisms promote germination and plant growth. Most studies attribute bacteria to biostimulatory, biofertilizer, and biocontrol skills (Santos et al., 2019). Some research already reports that bacteria can positively modulate the mobilization of seed reserves during germination. In this study, the changes in compositional bacteria structure do not seem to interfere with this seed-stored mobilization, maybe due to the remaining presence of key taxon after disinfection. On the other hand, if the number of bacteria/ng of DNA (detected via real-time PCR) has not been altered by disinfection, the mobilization of reserves will not be altered either.

After the loss of several germination assays due to the action of the pathogenic fungus *Penicillium* sp. (data not shown from our group for *Z. mays* variety SHS5050), we decided to explore the role of the bacterial microbiota in protecting the seed (maize of the variety DKB177). Results of this test showed that the partial removal of the microbiota by the hypochlorite rendered the seed more susceptible to the seed-borne fungus *Penicillium* sp, drastically reducing the germination and growth of maize. On the other hand, seeds that were not disinfected and inoculated with the fungus developed typically, as well as the control seeds without fungus challenger. This finding was confirmed by SEM of germinated roots, with dense fungal colonization in the disinfected-inoculated treatment, while little or no hypha was observed in the treatment non-disinfected-inoculated. Other studies have also shown that seed-bacteria control fungal diseases (Verma et al. 2017; Khalaf and Raizada 2018; Verma et al., 2018; Verma and White, 2018; White et al. 2017).

We correlated biocontrol results with maize sequencing and observed that the bacteriome acted as a “barrier” against phytopathogens by inhibiting the proliferation of fungi that deteriorate the seeds, also determining the bacterial profile of the root and the growth parameters of the maize seedlings. For this, they had to compete for nutrients and niches, induce plant resistance or produce antifungal and antibacterial metabolites (antibiotics, bacteriocins, lytic enzymes, and volatile compounds) (Verma et al., 2019). The candidates’ bacteria consortium responsible for microbial “barrier” are f_Enterobacteriaceae_922761, *Azospirillum,* and *Acinetobacter*. With disinfection, the abundance of these genera was reduced, changing the root bacteriome, making the seed susceptible to *Penicillium* sp. that harming the germination and seedling growth of maize.

Possible functional abilities of the featured gender have been established in the literature. Recent studies indicate that endophytic strains of *Enterobacter* (classified in the family f_Enterobacteriaceae_922761) can stimulate germination and plant growth (Panigrahi et al., 2019, Vitorino et al., 2019). The underlying mechanisms involve phytohormones (indole-acetic acid-IAA) (Verma et al., 2017, Srisuk et al., 2018, Panigrahi et al., 2019), siderophores (Maleki et al., 2018, Panigrahi et al., 2019) and phosphate solubilization (Verma et al., 2017, Panigrahi et al., 2019, Luduena et al., 2018); and that seeds not treated with these bacteria are susceptible to degradation by fungal phytopathogens, such as *Penicillium*, *Fusarium* and others (Sandhya et al., 2017, Verma et al., 2017, Vitorino et al., 2019). Enterobacteria were able to inhibit the growth of *Aspergillus flavus* and seven other fungal pathogens through volatile compounds produced (Gong et al., 2019).

The *Azospirillum* genus comprises bacteria widely studied and used in agriculture (Santos et al., 2019). Its inoculation in plants promotes growth through different mechanisms, such as biological nitrogen fixation, production of phytohormones (such as IAA, gibberellins), and siderophores (Fukami et al., 2017, López-Reyes et al., 2017). This genus has been attributed to the ability to reduce the incidence of fungal diseases (*Alternaria*, *Bipolaris,* and *Fusarium*) of maize (López-Reyes et al., 2017) through the induction of defense genes (Fukami et al., 2017, Fukami et al., 2018).

Other studies have identified in *Acinetobacter* bacteria the ability to produce IAA, siderophores, solubilize phosphate and zinc, fix nitrogen and promote plant growth (Gandhi and Muralidharan, 2016, Kang et al., 2016, Patel et al., 2017); in addition to acting on the control of fungi associated with seeds (*Fusarium* and *Alternaria*) (Medina-de la Rosa et al., 2016) through chitinases they produce (Krithika and Chellaram, 2016).

## Conclusion

We concluded that the structure of the seed-borne bacteria community is drastically shaped (mainly taxon relative abundance) by the germination process of maize that ultimately influences germination and seedling growth. Additionally, the removal of certain bacteria taxa by chemical seed-disinfection suppress natural seed-borne barrier protection of maize seedlings from fungal pathogens. Understanding the successional community balance during the seed germination and critical community members of the microbial network and physiological process will open up new ways for the formulation of inoculants to boost crop productivity and crop protection. Although the strategy of creating seed microbiome-based inoculants has not yet been put into practice, it represents the future of agriculture.

## Supporting information

Distribution of reads per sample of the 5-h water embedded no germinated seed.

Permanova of the seed bacteriome.

Distribution of reads per sample of the 5-d emerged root from germinated seeds.

Permanova of the root bacteriome.

Differential abundance of roots from disinfected versus non-disinfected seeds.

. Differences in protein, glucose, triglyceride, reducing sugar, and alpha-amylase activity from different stages of maize germination

## Acknowledgments

FAPERJ grant n° E-26/203.003/2017, CNPq grant n° 314263/2018-7, Newton Fund grant BB/N013476/1 “Understanding and Exploiting Biological Nitrogen Fixation for Improvement of Brazilian Agriculture,” co-funded by the Biotechnology and Biological Sciences Research Council (BBSRC) and the Conselho Nacional das Fundações Estaduais de Amparo à Pesquisa (CONFAP) and FINEP-PLURICANA financially supported this study. This study is part of the Ph.D. of the first author (LFS), who is grateful for the fellowship conceded by CAPES.

## Conflict of interest

No conflict of interest declared.

## Appendices caption

**Fig. A.1.**
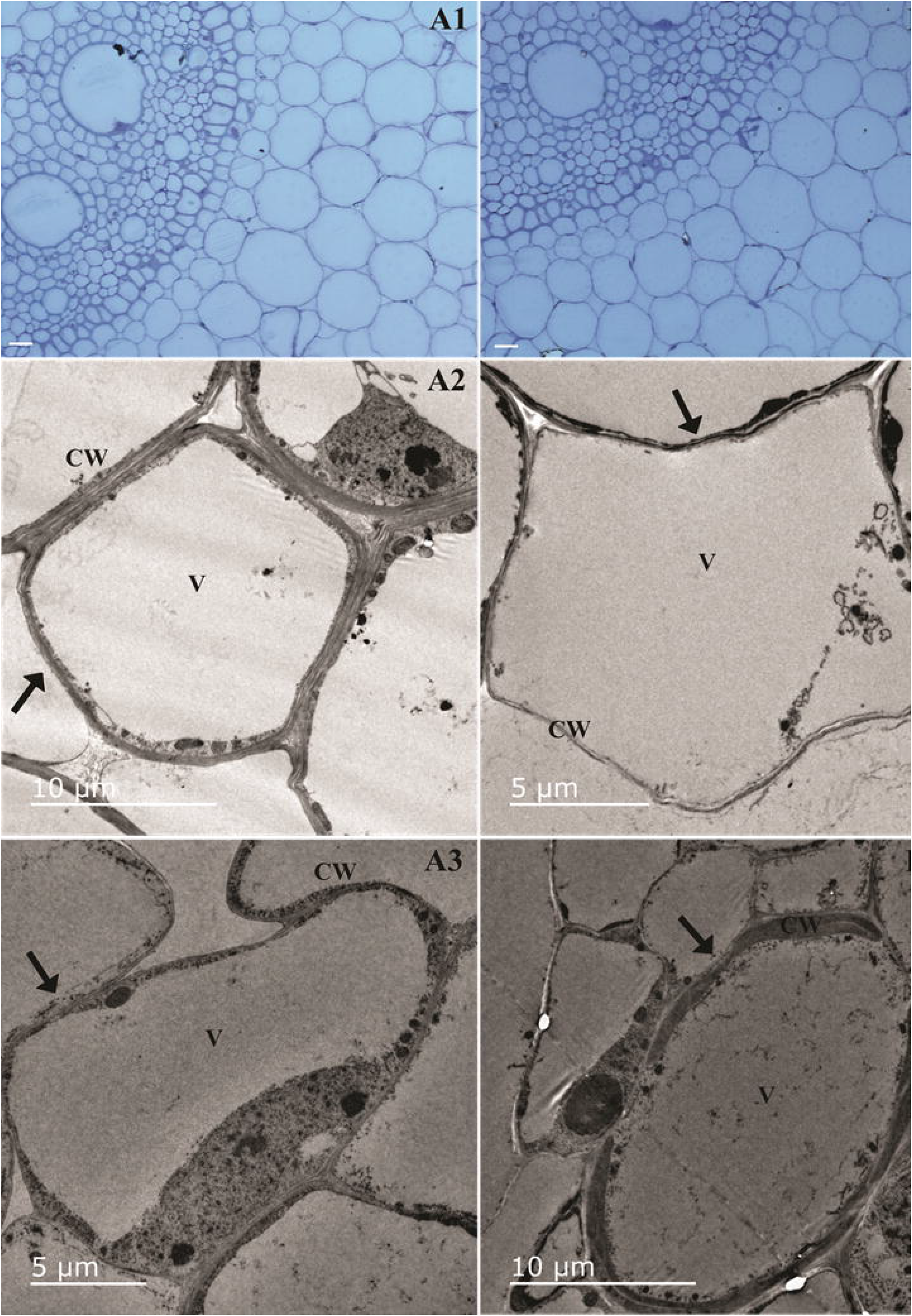
Root tip cross-sections of non-disinfected (A) and disinfected (B) maize seeds viewed by light microscopy (A1 and B1) and transmission electron microscopy (A2, A3, B2, and B3). Black arrows indicate cell wall shape. Plant cell organelles: cell wall (CW) and vacuole (V). Bars represent the following scales: panel A1 and B1: 20 μm; A2 and B3: 10 μm; B2 and A3: 5 μm.

**Fig. A.2.**
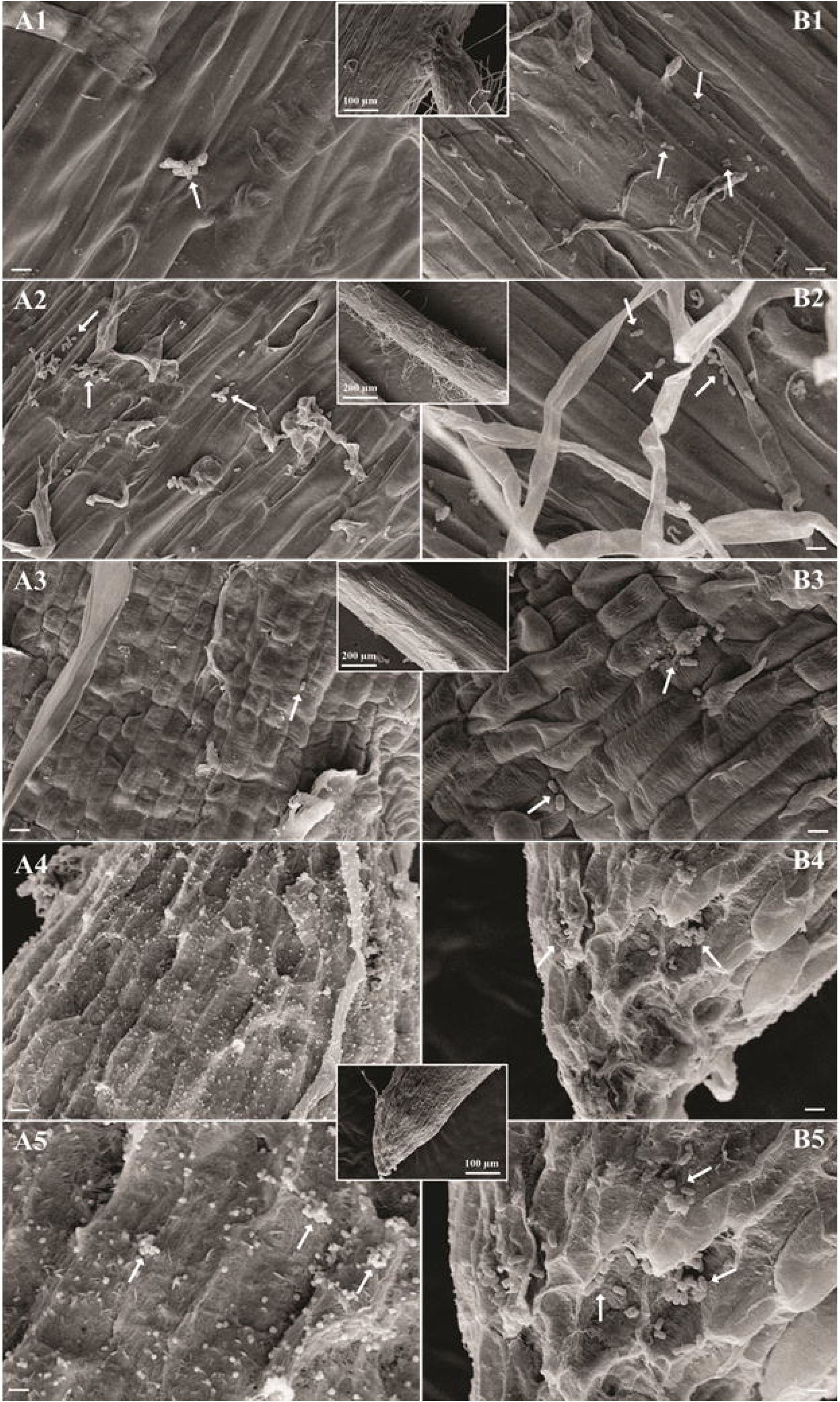
Bacterial colonization of maize root visualized by scanning electron microscopy (SEM). Maize seeds non-disinfected (A) and disinfected (B). Root regions: lateral root (A1 and B1), root hair (A2 and B2), zone of elongation (A3 and B3), root cap (A4, A5, B4, and B5). Bacteria and yeast are indicated by arrows. Bars represent the following scales: panel A1 and A5: 20 μm; A2-A4 and B1-B5: 20 μm.

## Highlights

- Seed-borne bacteria is an essential source of the plant microbiome.
- Maize seed germination is a secondary ecological succession for seed-borne microbiota.
- Seed-bacteriome modulates the abundance of taxa in germinated maize seeds.
- Seed-borne bacteria influence the germination and growth of maize.
- Seed microbiome acts as a “bacterial barrier” against phytopathogens.
- Maize seed-bacteria community var DKB177 protect against *Penicillium* sp. Seed-borne fungus.

## Graphical Abstract

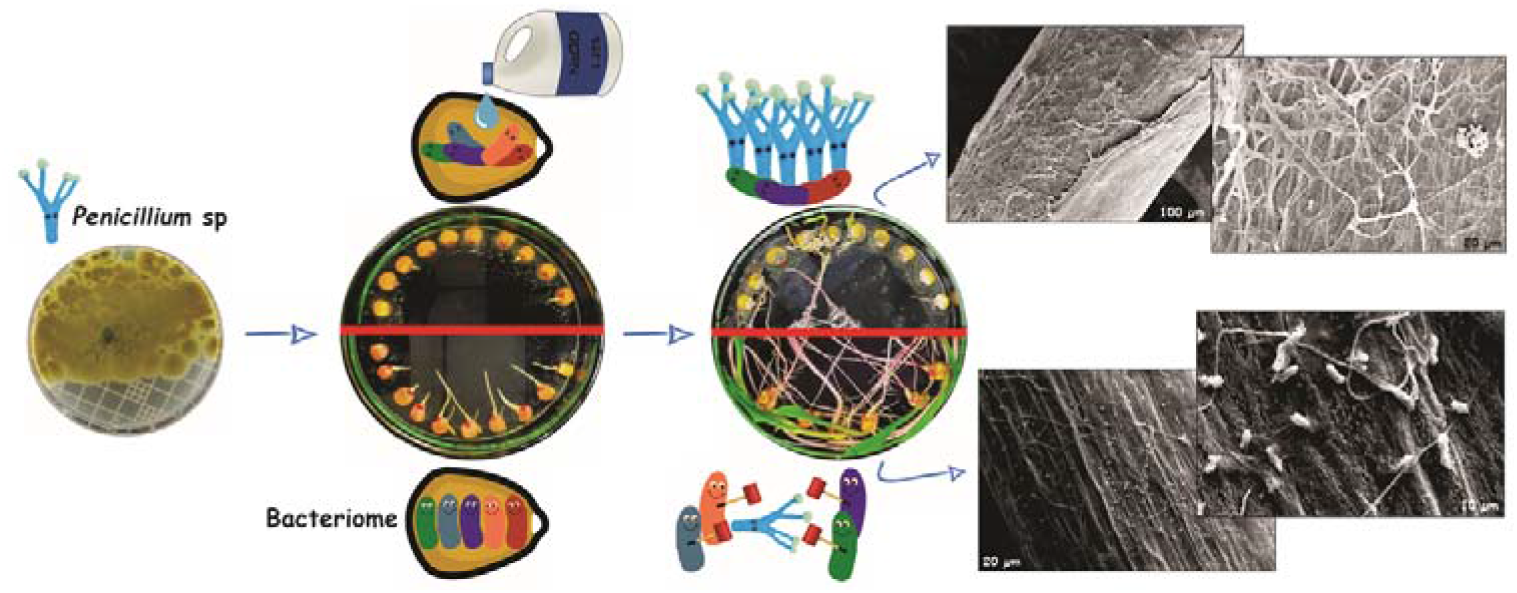

## Notes

### Competing Interest Statement

The authors have declared no competing interest.

