## Supplementary material for "Altered bacteria community dominance reduces tolerance to resident fungus and seed to seedling growth performance in maize (*Zea mays* L. var. DBK 177)": Distribution of reads per sample of the 5-h water embedded no germinated seed.

Supplementary Table 1. Distribution of reads per sample of the 5-h water embedded no germinated seed.

| **X.SampleID** | **Genotype** | **Extraction** | **Disinfection** | **Treatment** | **TotalReads** | **Coverage** |
| --- | --- | --- | --- | --- | --- | --- |
| **O34** | DKB177 | CTAB | No | S_DKB177_ND_CTAB | 106 | 0.6509434 |
| **O33** | DKB177 | CTAB | Yes | S_DKB177_D_CTAB | 92 | 0.4239130 |
| **O32** | DKB177 | CTAB | Yes | S_DKB177_D_CTAB | 183 | 0.7868852 |
| **O36** | DKB177 | CTAB | No | S_DKB177_ND_CTAB | 61 | 0.4262295 |
| **O35** | DKB177 | CTAB | No | S_DKB177_ND_CTAB | 62 | 0.4516129 |
| **O31** | DKB177 | CTAB | Yes | S_DKB177_D_CTAB | 83 | 0.4216867 |
