## Supplementary material for "Altered bacteria community dominance reduces tolerance to resident fungus and seed to seedling growth performance in maize (*Zea mays* L. var. DBK 177)": Permanova of the seed bacteriome.

Table 2: Permanova of the seed bacteriome.

|  | Df | SumsOfSqs | MeanSqs | F.Model | R2 | Pr(>F) |
| --- | --- | --- | --- | --- | --- | --- |
| Disinfection | 1 | 0.282091 | 0.2820910 | 0.9659997 | 0.1945227 | 0.6 |
| Residuals | 4 | 1.168079 | 0.2920197 | NA | 0.8054773 | NA |
| Total | 5 | 1.450170 | NA | NA | 1.0000000 | NA |
