## Supplementary material for "Altered bacteria community dominance reduces tolerance to resident fungus and seed to seedling growth performance in maize (*Zea mays* L. var. DBK 177)": Distribution of reads per sample of the 5-d emerged root from germinated seeds.

Supplementary Table 3. Distribution of reads per sample of the 5-d emerged root from germinated seeds.

| **X.SampleID** | **Genotype** | **Extraction** | **Disinfection** | **Treatment** | **TotalReads** | **Coverage** |
| --- | --- | --- | --- | --- | --- | --- |
| **O18** | DKB177 | CTAB | No | R_DKB177_ND_CTAB | 954 | 0.9245283 |
| **O24** | DKB177 | CTAB | No | R_DKB177_ND_CTAB | 126 | 0.7222222 |
| **O19** | DKB177 | CTAB | Yes | R_DKB177_D_CTAB | 274 | 0.8686131 |
| **O16** | DKB177 | CTAB | No | R_DKB177_ND_CTAB | 373 | 0.8176944 |
| **O15** | DKB177 | CTAB | Yes | R_DKB177_D_CTAB | 409 | 0.8459658 |
| **O14** | DKB177 | CTAB | Yes | R_DKB177_D_CTAB | 492 | 0.8231707 |
| **O21** | DKB177 | CTAB | Yes | R_DKB177_D_CTAB | 582 | 0.9123711 |
| **O20** | DKB177 | CTAB | Yes | R_DKB177_D_CTAB | 17979 | 0.9987764 |
| **O13** | DKB177 | CTAB | Yes | R_DKB177_D_CTAB | 435 | 0.8459770 |
| **O17** | DKB177 | CTAB | No | R_DKB177_ND_CTAB | 493 | 0.8194726 |
| **O23** | DKB177 | CTAB | No | R_DKB177_ND_CTAB | 417 | 0.8201439 |
| **O22** | DKB177 | CTAB | No | R_DKB177_ND_CTAB | 389 | 0.8868895 |
