## Supplementary material for "Altered bacteria community dominance reduces tolerance to resident fungus and seed to seedling growth performance in maize (*Zea mays* L. var. DBK 177)": Permanova of the root bacteriome.

Suplementary Table 4: Permanova of the root bacteriome.

|  | Df | SumsOfSqs | MeanSqs | F.Model | R2 | Pr(>F) |
| --- | --- | --- | --- | --- | --- | --- |
| Disinfection | 1 | 0.3394742 | 0.3394742 | 0.9995477 | 0.0908717 | 0.384 |
| Residuals | 10 | 3.3962776 | 0.3396278 | NA | 0.9091283 | NA |
| Total | 11 | 3.7357517 | NA | NA | 1.0000000 | NA |
