## Supplementary material for "Altered bacteria community dominance reduces tolerance to resident fungus and seed to seedling growth performance in maize (*Zea mays* L. var. DBK 177)": Differential abundance of roots from disinfected versus non-disinfected seeds.

| rab.all | rab.win.R_  DKB177_D | rab.win.R_  DKB177_ND | effect | overlap | we.ep | Family | Genus | Species |
| --- | --- | --- | --- | --- | --- | --- | --- | --- |
| 3.987.430 | 58.208.770 | 177.209 | -10.068.706 | 8.072.996 | 312.351 | f__Moraxellaceae | g__*Acinetobacter* | NA^*^ |
| 4.928.825 | 61.503.817 | 437.454 | -5.952.440 | 24.739.603 | 1.441.941 | f__Moraxellaceae | g__*Acinetobacter* | s__*rhizosphaerae* |
| 1.314.203 | -3.024.789 | 314.261 | 6.797.912 | 18.701.328 | 1.500.234 | f__Brucellaceae | g__*Ochrobactrum* | NA^*^ |

Supplementary Table 5. Differential abundance of roots from disinfected versus non-disinfected seeds.

^*^NA = unassigned species
