## Supplementary material for "Altered bacteria community dominance reduces tolerance to resident fungus and seed to seedling growth performance in maize (*Zea mays* L. var. DBK 177)": . Differences in protein, glucose, triglyceride, reducing sugar, and alpha-amylase activity from different stages of maize germination

Supplementary Table 6. Differences in protein, glucose, triglyceride, reducing sugar, and alpha-amylase activity from different stages of maize germination (seed, embryo axis, and root).

| **Treatments** | | |
| --- | --- | --- |
|  | **NDS** | **DS** |

***Protein (ug/ul)***

| Seed | 0.2805^b^ | 0.1494^b^ |
| --- | --- | --- |
| Embryonic axis | 0.4385^b^ | 0.2706^b^ |
| Root | 1.7384^a^ | 1.3464^a^ |

***Glucose (mg/dl)***

| Seed | 0.0087^b^ | 0.0099^b^ |
| --- | --- | --- |
| Embryonic axis | 0.0213^b^ | 0.0189^b^ |
| Root | 0.0664^a^ | 0.0708^a^ |

***Triglyceride (mg/dl)***

| Seed | 0.0079^a^ | 0.0034^b^ |
| --- | --- | --- |
| Embryonic axis | 0.0082^a^ | 0.0069^a^ |
| Root | 0.0090^a^ | 0.0087^a^ |

***Reducing sugar(ug/ul)***

| Seed | 0.1471^b^ | 0.1390^b^ |
| --- | --- | --- |
| Embryonic axis | 0.1738^b^ | 0.1510^b^ |
| Root | 0.2765^a^ | 0.2756^a^ |

***Alpha-Amylase (U/g)***

| Seed | 14.8333^c^ | 8.0000^c^ |
| --- | --- | --- |
| Embryonic axis | 35.6667^b^ | 24.0000^b^ |
| Root | 46.9167^a^ | 47.6667^a^ |

Maize seeds non-disinfected (NDS) and disinfected (DS). Different letters in the same column indicate significant differences between stages, according to the Tukey test (p ≤ 0.05).
